# StarTrace: A Multiplex Organoid Avatar Drug Testing Platform for Personalized Medicine

**DOI:** 10.1101/2025.02.12.637574

**Authors:** Sana Khalili, Shrey Patel, Nidhi Patel, Victoria Moy, Sawyer Lyons, Emma Gray, Riley Brents, Carolyn E. Banister, Sidney E. Morrison, Phillip J. Buckhaults

## Abstract

The goal of precision medicine is to improve clinical outcomes of cancer patients by choosing treatments most likely to work. One idea is to match tumor response to cancer-causing somatic mutations, but this strategy still faces limitations in colorectal cancer due to poorly understood genetic influences on drug resistance. We describe here a simple direct drug sensitivity assay platform applied to mixtures of patient-derived organoid avatars as a practical solution for choosing therapy, sidestepping the need for exhaustive knowledge of drug-genetic interactions. This approach rank orders individual patient’ organoid avatars’ responses to various drugs to be used to guide treatment choices. The platform multiplexes organoids using a clonal barcoding method, called StarTrace, which simultaneously tests pools of multiple patients’ organoid avatars for sensitivity or resistance to small molecule inhibitors. We utilized both quantitative real-time PCR-based and single-molecule sequencing assays to track the relative Darwinian fitness of each barcoded organoid within the pool. StarTrace offers a rapid, cost effective and sensitive testing platform that could be useful for either preclinical drug development or tailoring personalized therapy.

## Main

The development of cancer treatments tailored to the needs of individual patients is a promising paradigm for oncology. Increased utilization of tumor sequencing illustrates the growing interest in using therapies that specifically target genetic variations in tumors ^1^. However, while genetic sequencing has advanced the identification of targetable mutations, only a small percentage of tumors have mutations known to be actionable, and even those that do may not always respond predictably to treatment. Many tumors, despite lacking clear genetic indicators, may still be highly sensitive to drugs that are not typically considered for those patients. The disconnect between somatic mutations and therapeutic response highlights the need for more versatile approaches to cancer treatment.

*In vitro* anticancer drug screens have traditionally been performed using 2D cancer cell lines, often using concentrations of drug that produce cell death within 2-3 days. However, these preclinical results often correlate poorly with animal models or with actual clinical outcomes ^2^. There is an unmet need for more relevant preclinical models that can be used at early stages of drug discovery such as high-throughput screening (HTS). Preclinical models should represent the recurring genetic themes present in real human cancers, enabling more relevant preclinical results and product development decisions. Patient-derived organoid avatars (PDOs) are sophisticated and highly relevant *in vitro* models. Organoids are self-organizing mammalian adult stem cells and are strong tools for *ex vivo* tissue morphogenesis and organogenesis simulations ^3^. Because cancer and normal organoids contain the array of germline and somatic mutations that influence drug response, this technology has the potential to bridge the gap between our basic science understanding of cancer genetics and pragmatic testing of new treatments for patients. By using PDOs, it is possible to capture the heterogeneity of human tumors, providing more accurate readout of potential treatment efficacy. Organoids can be used alongside traditional methods such as cell-line and xenograft-based drug research and have the advantage of enabling individualized therapy design ^4^. Here we show a new methodology, called StarTrace, for high-throughput *in vitro* drug screening of a pool of genetically barcoded colorectal cancer organoid avatars. Drug effect is measured over long periods of time in terms of Darwinian fitness using both next-generation sequencing (NGS) and qPCR to quantify each barcode in the mixture. This method is highly sensitive and more relevant to patient outcomes than are typical short-term cell death assays.

## Results

### High-Throughput Barcode Deconvolution Using Next-Generation Sequencing

Molecular barcoding, involving the integration of short, non-coding DNA sequences into the genomes of cell populations via viral transduction has been used previously to monitor the invisible sub-clones within the diverse tumor cell populations, providing insights into tumor development and progression ^5^ ^6^ ^7^ ^8^ ^9^ ^10^ ^11^ ^12^ ^13^ ^14^ ^15^ ^16^. We developed an NGS-based method that enables to track each patient-derived organoids (PDOs) present in a mixture in response to treatment with various cancer drugs. This method uses oxford nanopore amplicon sequencing to track the frequency of each unique barcode during extended culturing of the mixture in various drug conditions. We isolated 72 randomly picked barcode clones from a high-complexity ClonTracer barcoding library (addgene #67267) ^5^ and generated infectious viral particles for tagging individual organoids (Fig. 1). Each barcode, comprising a 30-base pair unique DNA sequence, was identified by Sanger sequencing (Supplementary Fig. 1). We selected barcode sequences sufficiently divergent from one another (Fig. 2a-b), to allow reliable identification of each barcode when mixtures are sequences by nanopore-platform amplicon sequencing. All organoids were characterized by tumor/normal whole exome sequencing (supplementary Data 1) and select cancer-causing somatic mutations ^17^ ^18^ ^19^ are shown in Table1.

**Fig. 1:**
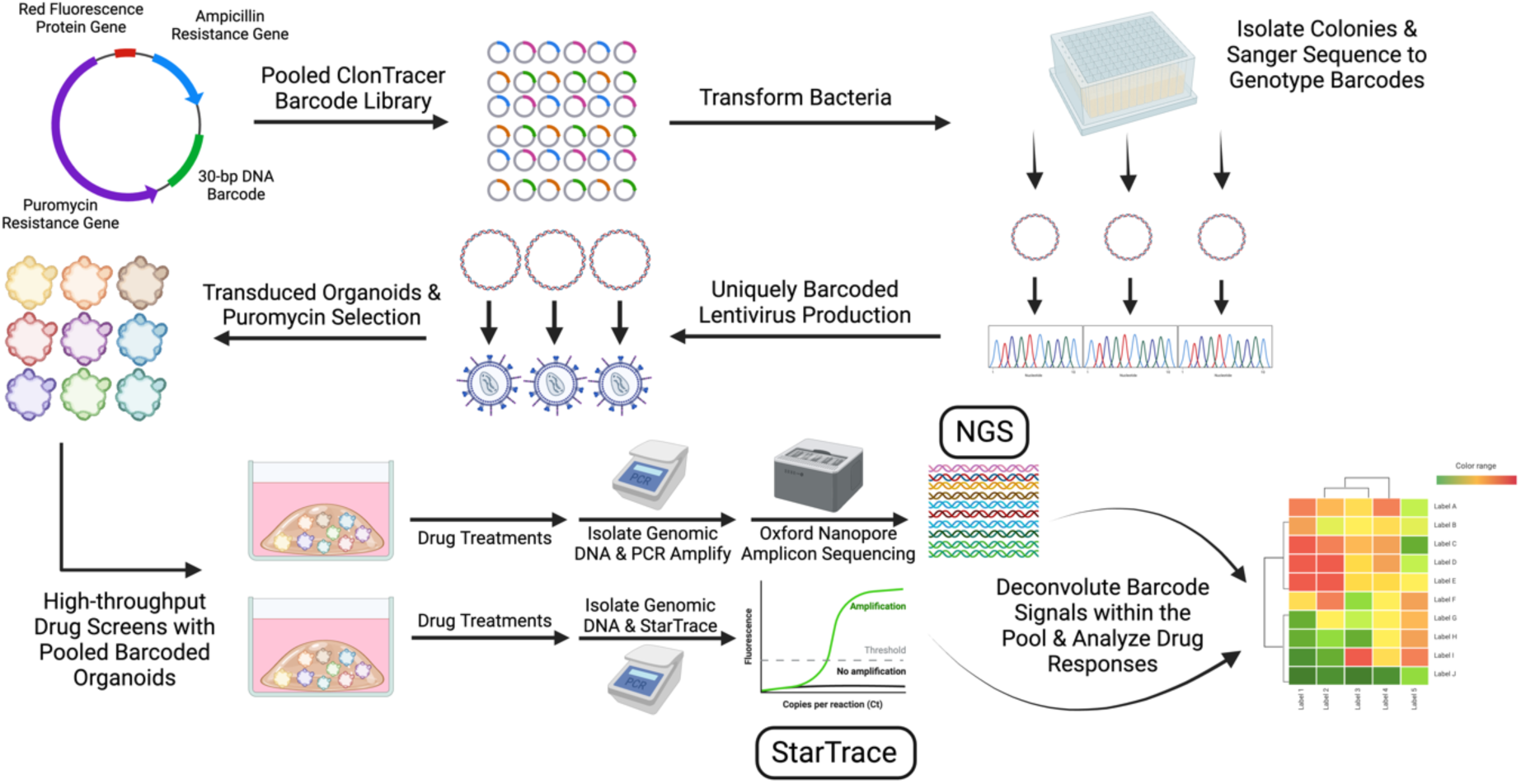
StarTrace Workflow. Isolation of individual DNA barcodes from the ClonTracer barcode virus library and workflow of barcoding and tracking PDOs within a mixture by both NGS and StarTrace PCR. Created in https://BioRender.com

**Fig. 2:**
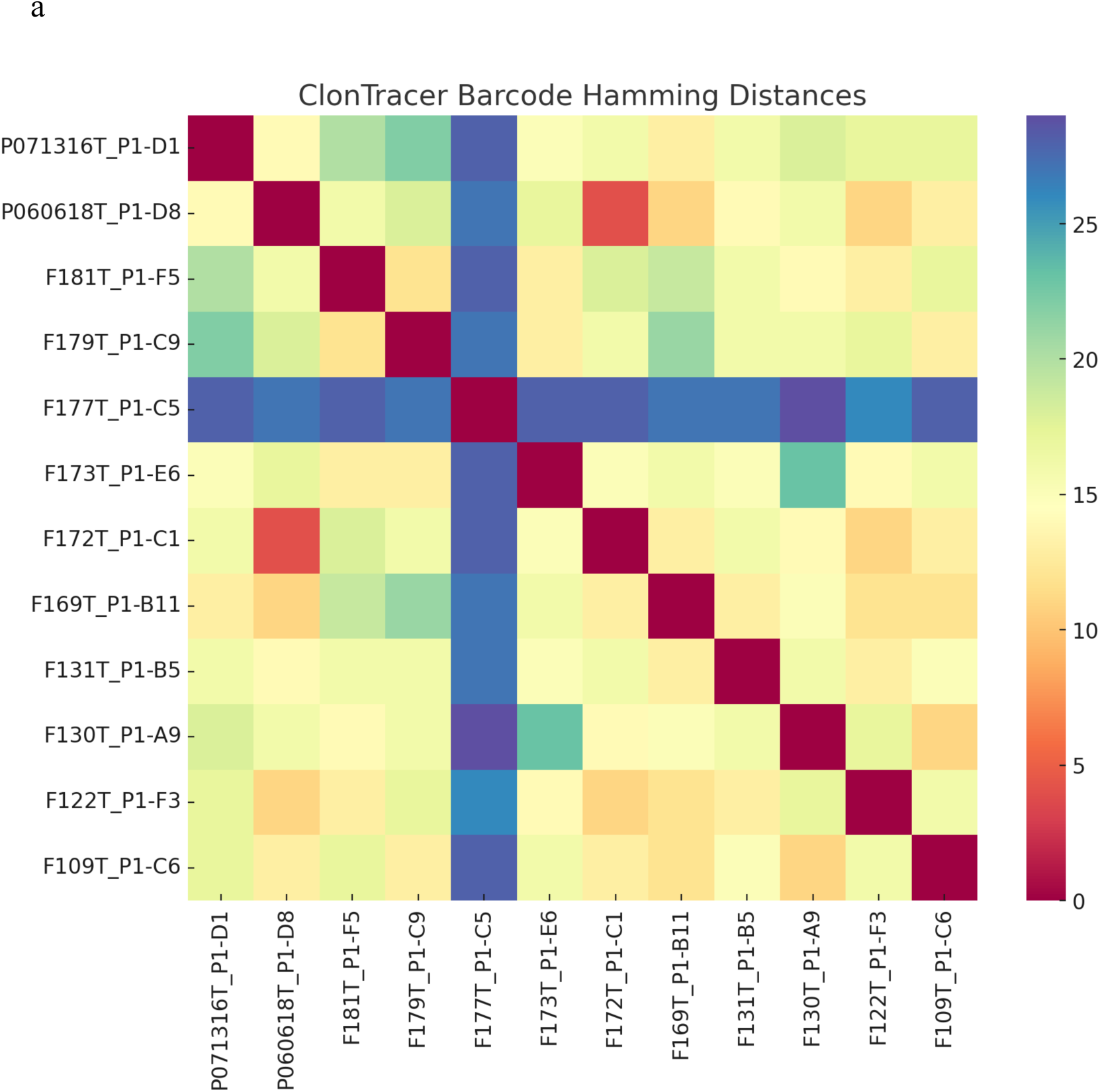
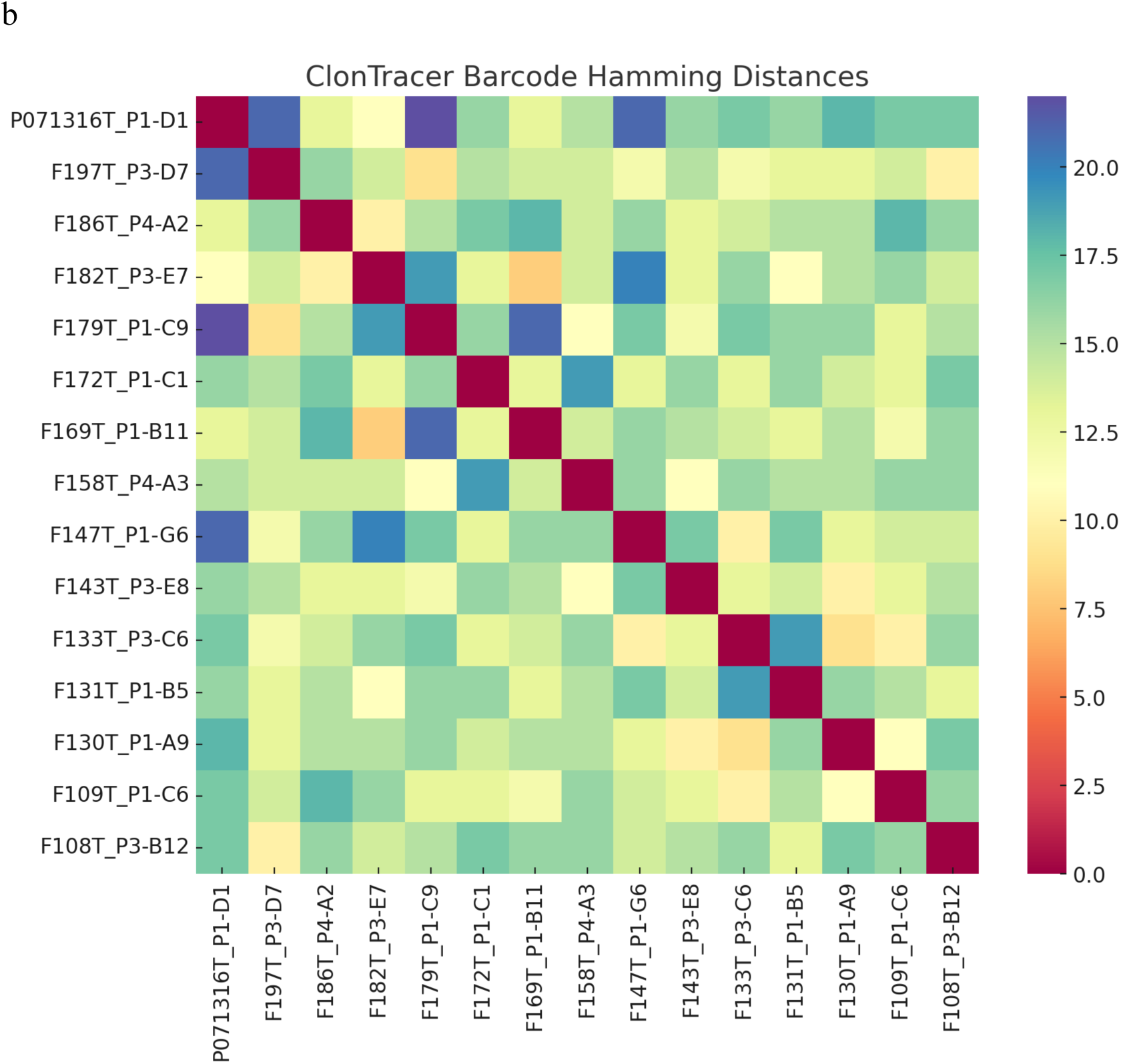
The uniqueness of DNA barcodes. **a,** Heatmap showing the pairwise hamming distances between 12 barcodes in the BC Pool-1. Each cell represents the hamming distance between two barcodes, red indicates low similarity, and blue indicates high similarity. **b,** Heatmap showing the pairwise hamming distances between 15 barcodes in the BC Pool-2. Each cell represents the hamming distance between two barcodes, red indicates low similarity, and blue indicates high similarity.

We transduced each individual PDO with a unique lentivirus clone containing a single DNA barcode and selected with puromycin. Each barcoded PDO contains lentiviral insertions containing a unique 30-bp barcode sequence flanked by lentiviral backbone sequences common to all barcoded organoids. These barcoded PDOs can be combined into bespoke organoid pools for different applications. We created two different custom pools to test cancer drug sensitivities. Pool-1 contained 12 barcoded PDOs and was treated with 5 different chemotherapy drugs, including nutlin-3a (MDM2 inhibitor), 5-Fluorouracil (5-FU), Mercaptopurine, Lapatinib, and Ibrutinib. Treatments were done with sub-lethal concentrations of each drug over a span of 28 days, with weekly sampling. Pool-2 contained 15 barcoded PDOs and was treated with nutlin-3a, LGK-974 (PORCN inhibitor), and Etoposide (TOPO isomerases inhibitor). Each sampling involved removing 50% of the organoid pool for qPCR and amplicon sequencing, while the remaining 50% was replated in fresh Matrigel for continued drug treatment. Throughout the process of DNA isolation, PCR, and sequencing, the size of each barcoded PDO population never dropped below a 1,000-fold bottleneck of 15,000 cells. Genomic DNA prepared from the time point samples of the pools was used for amplicon sequencing of the barcode locus on the Oxford Nanopore platform. We analyzed the FASTQ files using fuzzy grep commands (UGREP) and enumerated the counts of all expected barcodes. We computed the change in frequencies of each barcode over time to compute relative Darwinian finesses of each condition compared to the no drug control (Fig. 3a-b). As expected, nutlin resistance associated with TP53 mutations, indicating the presence of both TP53-Wildtype (TP53-WT) and TP53-Mutant (TP53-MT) organoids. Overall, 13 out of 20 PDOs were resistant to nutlin (Fig. 3a-b, Fig. 4, and Fig. 6), with 12 of these 13 nutlin-resistant PDOs found by whole exome sequencing (WES) to harbor inactivating somatic mutations in the TP53 gene (Table 1). One nutlin-resistant PDO, F143T_P3-E8 had no detectable TP53 somatic mutation. This nutlin resistance may result from either an undetected TP53 mutation or MDM2 amplification ^20^ (Table 1, Fig. 4 and Fig. 6).

**Table. 1:** Relationship between drug response and select somatic mutations. Somatic mutations in PDOs detected by WES.

| Organoid | Nutlin | Ibrutinib | Lapatinib | TKI | BRAF | ERBB3 | KRAS | TP53 |
| --- | --- | --- | --- | --- | --- | --- | --- | --- |
| F108T | R |  |  |  |  |  |  | p.Ser215Asn |
| F109T | R | R | R | R | p.Val600Glu |  |  | p.Arg234Cys |
| F122T | R |  |  |  |  |  |  | p.Asn92fs |
| F133T | R |  |  |  |  |  |  | p.Arg273His |
| F143T | R |  |  |  |  |  |  |  |
| F169T | R | ? | R | R |  |  | p.Gly12Asp | p.Gly245Ser |
| F172T | R |  |  |  |  |  |  | p.Arg248Gln |
| F179T | R |  |  |  |  |  |  | p.Arg136His |
| F181T | R | R | R | R |  |  | p.Gly12Asp | p.Ser127_Gln128dup |
| F182T | R |  |  |  |  |  |  | p.Cys176Tyr |
| F197T | R |  |  |  |  |  |  | p.Arg248Trp |
| P060618T | R | S | R | ? |  |  | p.Gly12Ala | p.Arg248Trp |
| P071316T | R | S | S | S |  | p.Ser1119Cys |  | p.Arg248Gln |
| F130T | S | R | R | R |  |  | p.Gly12Val |  |
| F131T | S | ? | R | R |  |  | p.Gly13Asp |  |
| F147T | S |  |  |  | p.Val600Glu |  |  | p.Val34fs<br>(Heterozygous) |
| F158T | S |  |  |  |  |  |  |  |
| F173T | S | S | ? | S |  | p.Val104Leu |  |  |
| F177T | S |  |  |  |  |  |  |  |
| F186T | S |  |  |  |  |  |  |  |

**Fig. 3:**
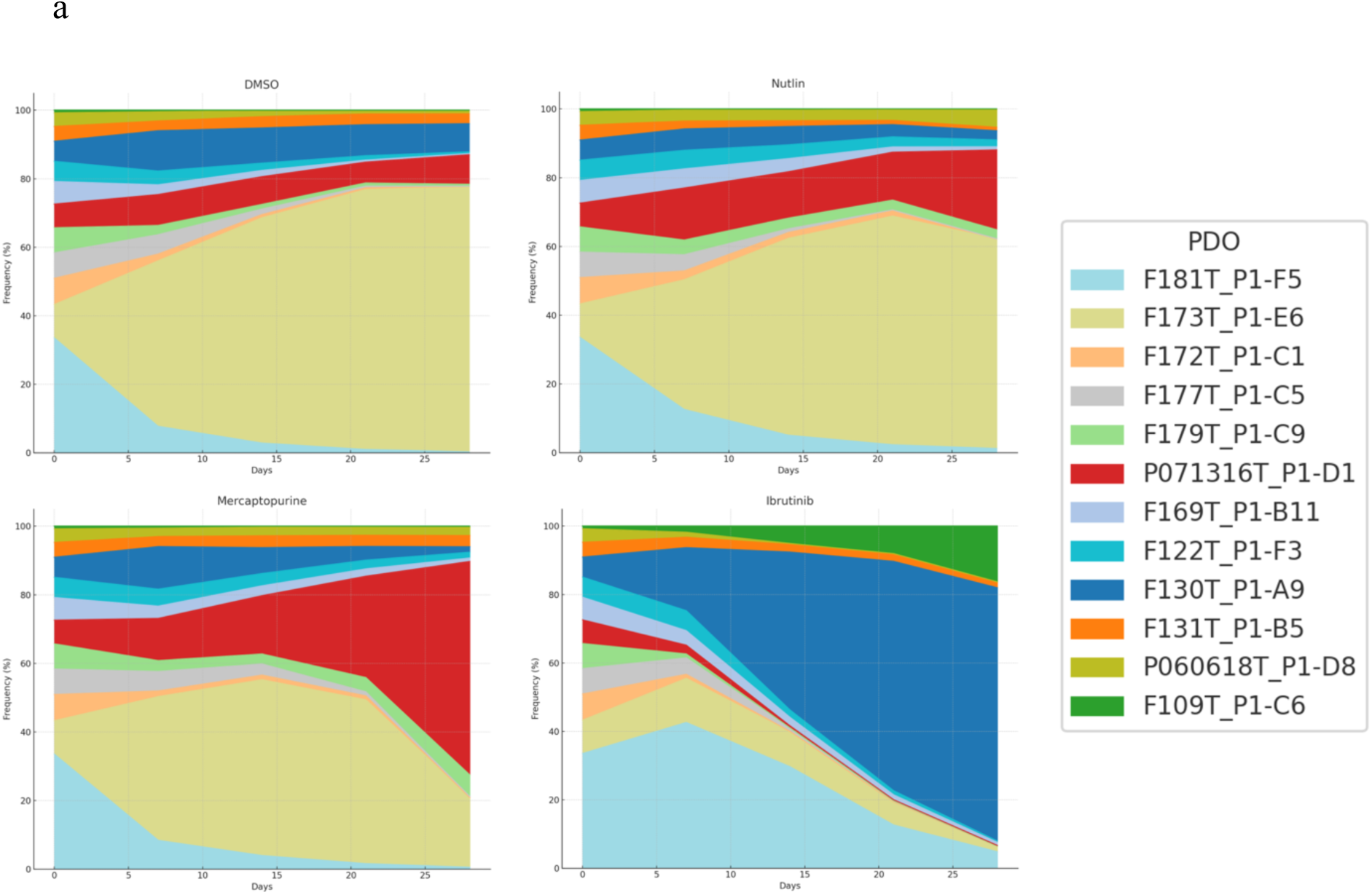
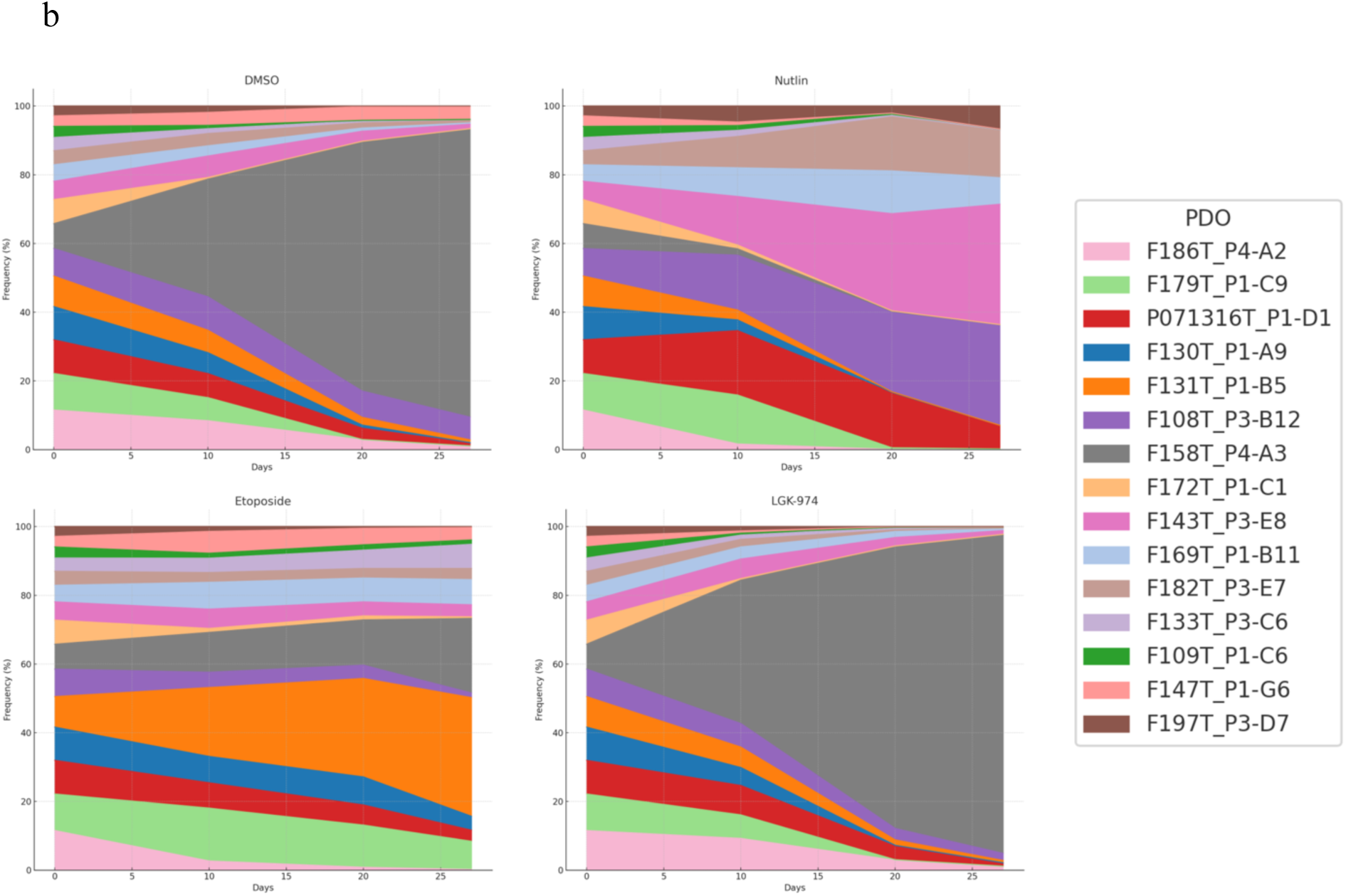
Change in frequencies of barcoded PDOs over time. a, Sand plots show the percentage survival of each barcoded organoids within the pool-1 over 28 days of treatment with drugs, only DMSO, Nutlin, Mercaptopurine, and Ibrutinib are shown. Each line represents an organoids’s survival trajectory, demonstrating how its proportion relative to the total pool changes over time. Some barcodes showing more rapid decline or resilience, suggesting differential susceptibility linked to genetic characteristics. All barcode frequencies were quantified by NGS sequencing of barcode amplicon. **b,** Sand plots show the percentage survival of each barcoded organoids within the pool-2 over 27 days of treatment with Nutlin, LGK-974, and Etoposide. Each line represents an organoids’s survival trajectory, demonstrating how its proportion relative to the total pool changes over time. Some barcodes showing more rapid decline or resilience, suggesting differential susceptibility linked to genetic characteristics. All barcode frequencies were quantified by NGS sequencing of barcode amplicon.

**Fig. 4:**
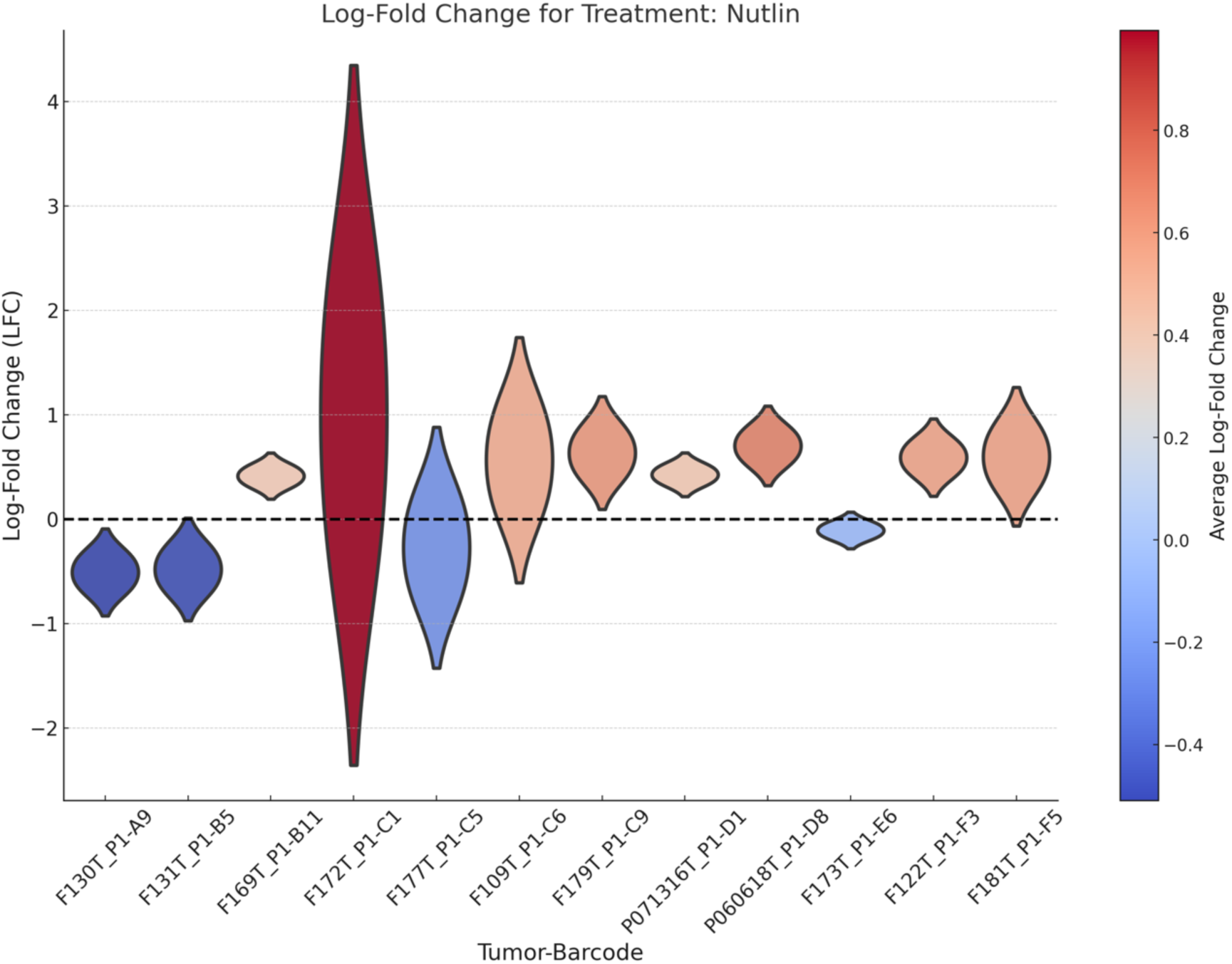
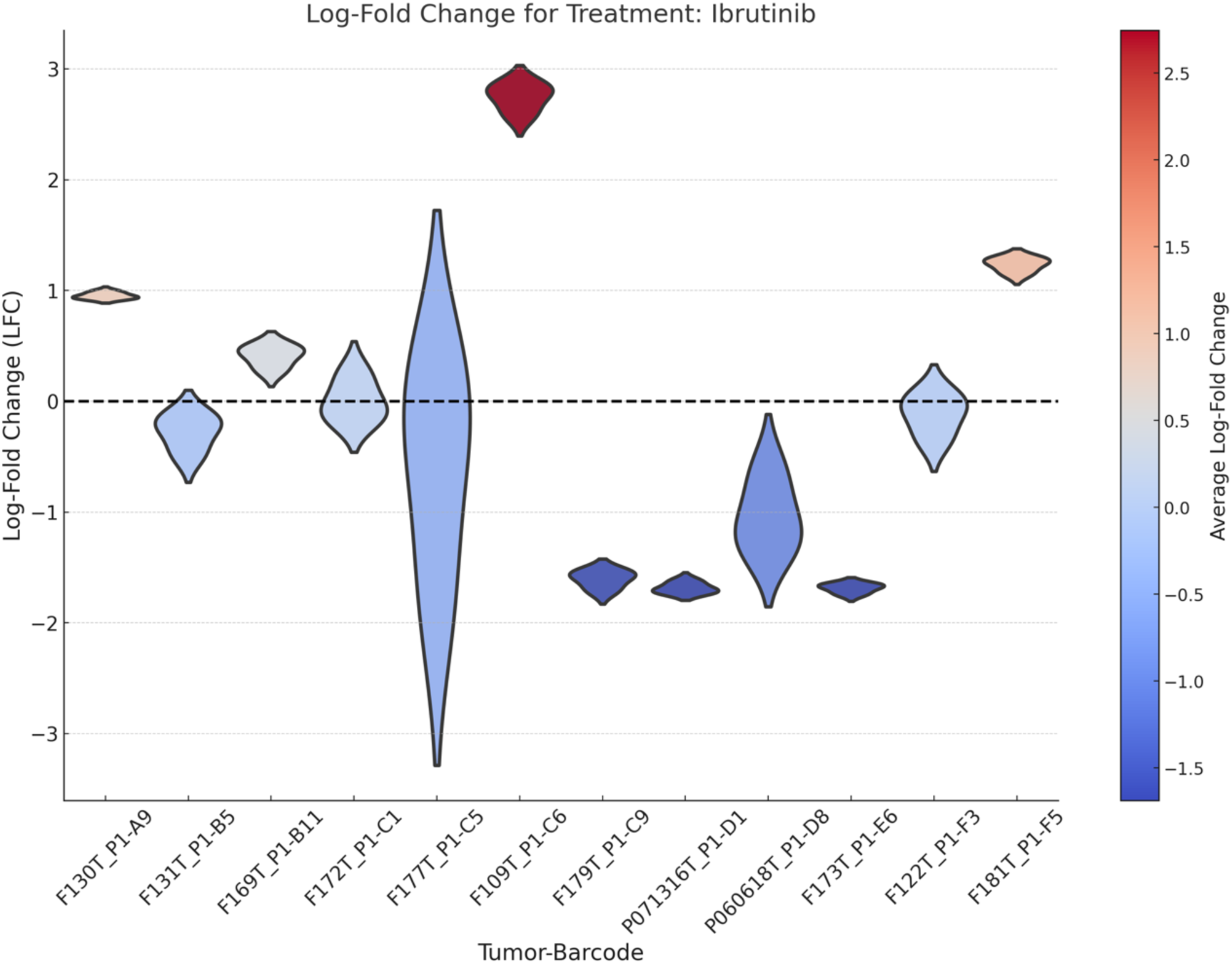
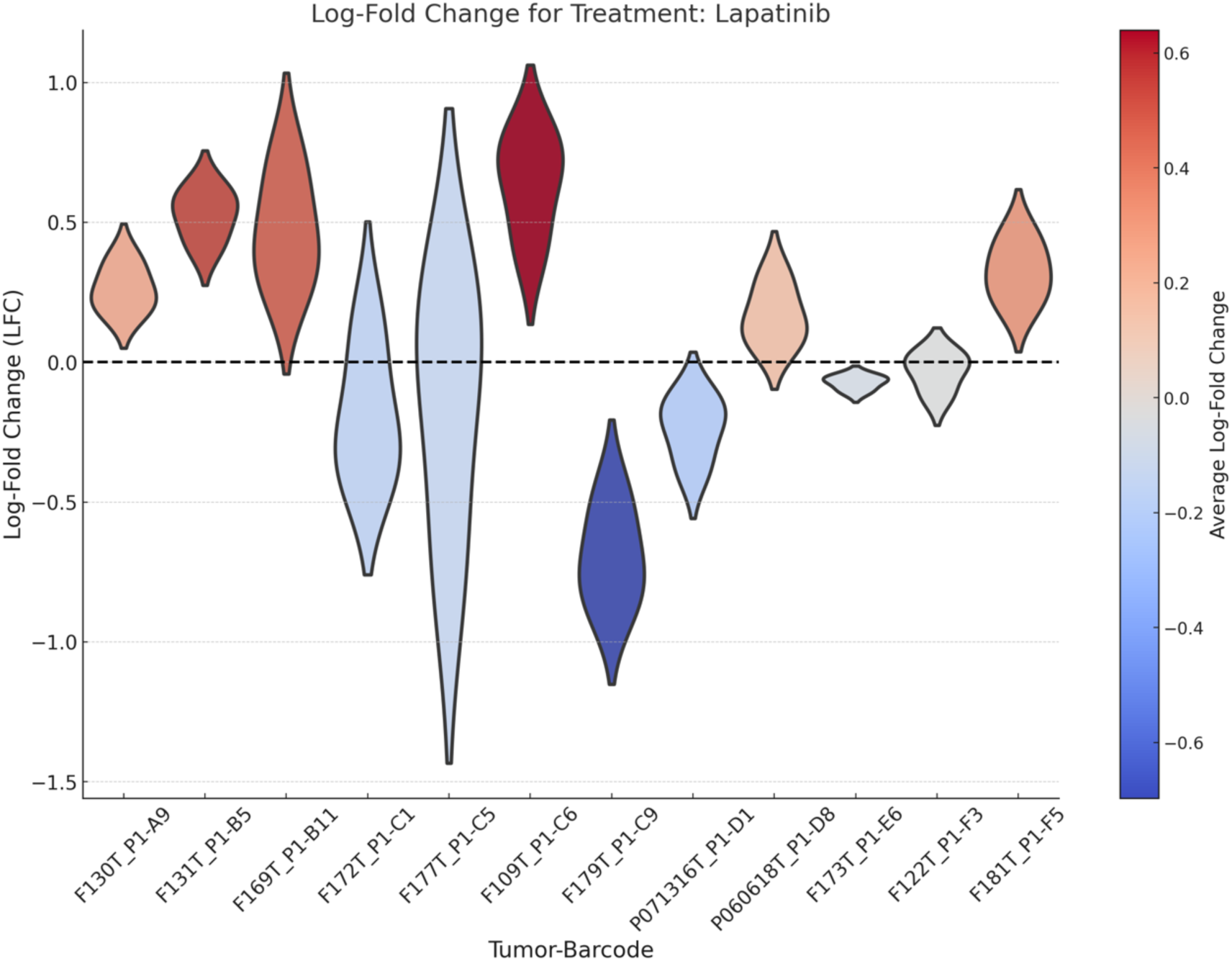
Drug responses of BC pool-1 PDOs over time, deconvoluted by NGS. Log-fold change in frequency of each barcoded PDOs within the pool-1, treated with different drugs for 28 days. Each violin represents the log-fold change in in a specific barcode relative to vehicle control.

Six organoids demonstrated resistance to tyrosine kinase inhibitors (TKIs), Ibrutinib and Lapatinib, and this correlated with the presence of BRAF and KRAS mutations ^21^. Two organoids (F173T-P1-E6 and P071316T_P1-D1) showed sensitivity to these inhibitors, which may be related to the presence of ERBB3 mutations in these tumors. (Tale 1, Fig. 4; Supplementary Fig. 2).

Ibrutinib is a tyrosine kinase inhibitor used to target Bruton’s tyrosine kinase (BTK) in hematological malignancies ^22^. The F173T PDO exhibited sensitivity to Ibrutinib, despite lacking any detectable BTK mutation. However, F173T carries a somatic mutation to ERBB3 (Val104Leu) which may heterodimerize with and trans activate other receptor tyrosine kinases ^23^. Figure 5a shows that the loss of mutant ERBB3 allele frequency with Ibrutinib treatment correlates well with the loss of the barcode for F173T. To determine if Ibrutinib sensitivity is a somatically acquired phenotype, we performed a drug response experiment using an unbarcoded mixture of F173T and patient-matched normal, F173N (Fig. 5b), followed by PCR sequencing of the mutant ERBB3 amplicon. The mutant ERBB3 allele was significantly depleted at high Ibrutinib concentrations. This result aligns with studies showing Ibrutinib inhibits ERBB receptor phosphorylation, leading to the suppression of key survival pathways such as PI3K/AKT and MAPK/ERK in solid tumors^24 25^. Overall, this result demonstrates the utility of functional testing in organoids to identify unexpected, yet relevant drug sensitivities.

**Fig. 5:**
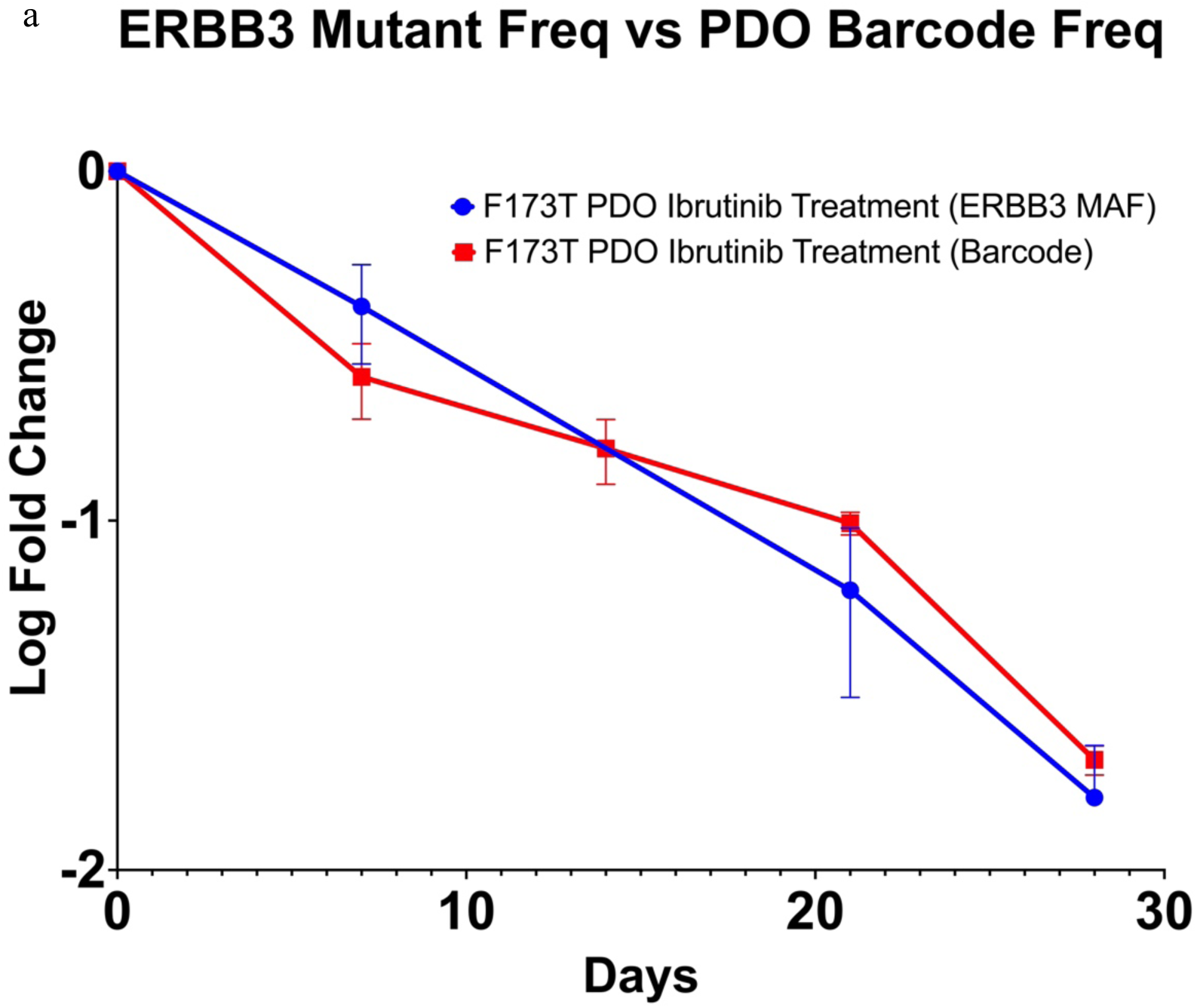
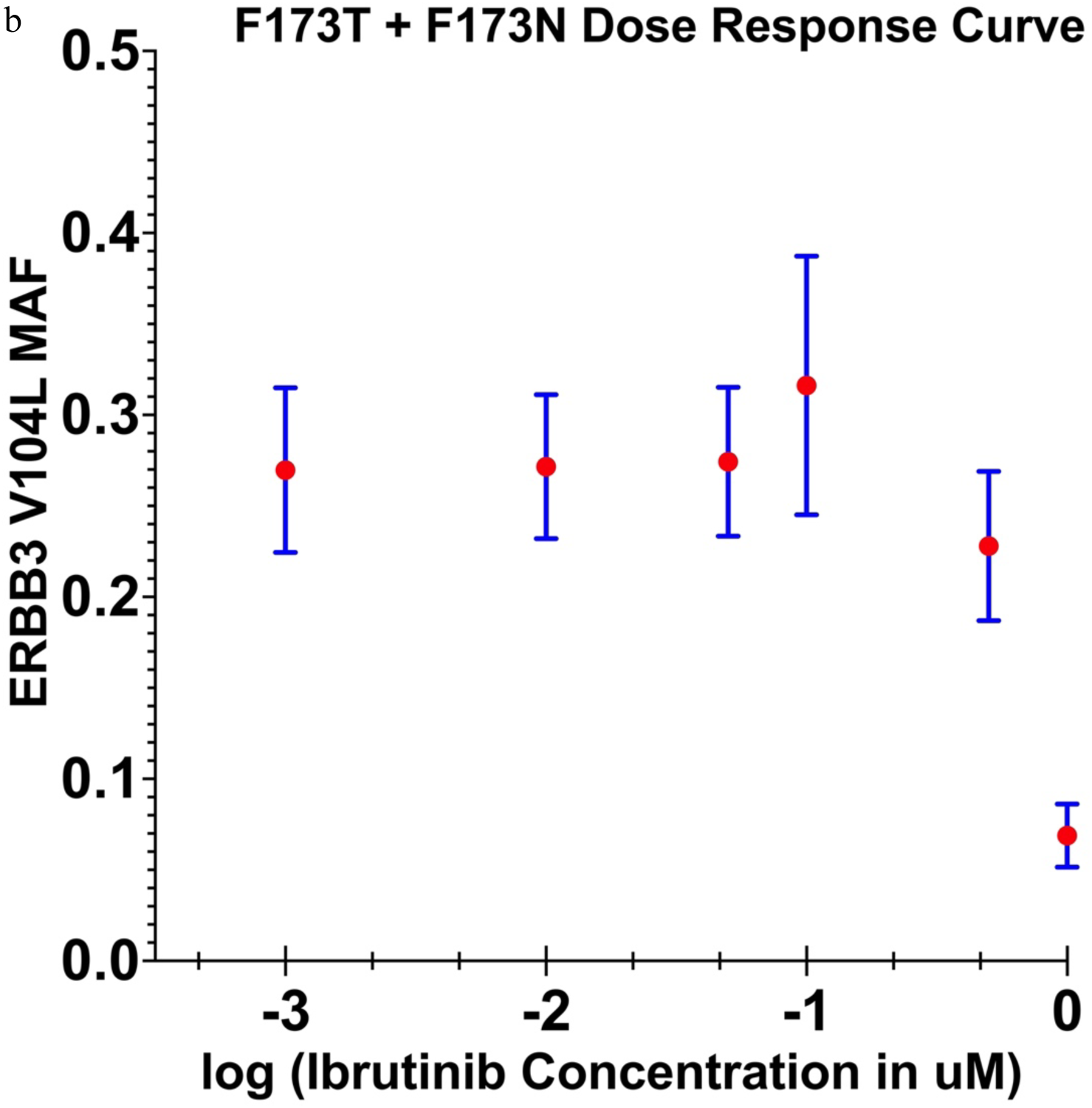
Ibrutinib response of an individual PDO with ERBB3 mutation, deconvoluted by NGS. **a,** Log fold change of both ERBB3 mutant frequency and barcoded F173T PDO frequency over time in presence of Ibrutinib. PDO frequency was determined by sequencing both the barcode amplicon and an ERBB3 amplicon flanking the Val104Leu mutant allele. **b,** Ibrutinib dose-response of F173T tumor and its matched normal F773N. The ERBB3 mutant allele frequency in a mixture of F173T (tumor) and F173N (normal) PDOs treated with Ibrutinib. PDO frequency was determined by sequencing an ERBB3 amplicon flanking the Val104Leu mutant allele. Data points represent the average of ERBB3 mutant allele frequency from three biological replicates.

Topoisomerase inhibitors are sometimes used in colon cancer treatment ^26^; however, outcomes remain difficult to predict due to the lack of reliable biomarkers of clinical response. PDOs showing both etoposide sensitivity and resistance are clearly identifiable (Fig. 3b and Fig. 6). Phenotyping cancers *ex vivo* may have practical utility for patients deciding on topoisomerase inhibitor therapy.

**Fig. 6.**
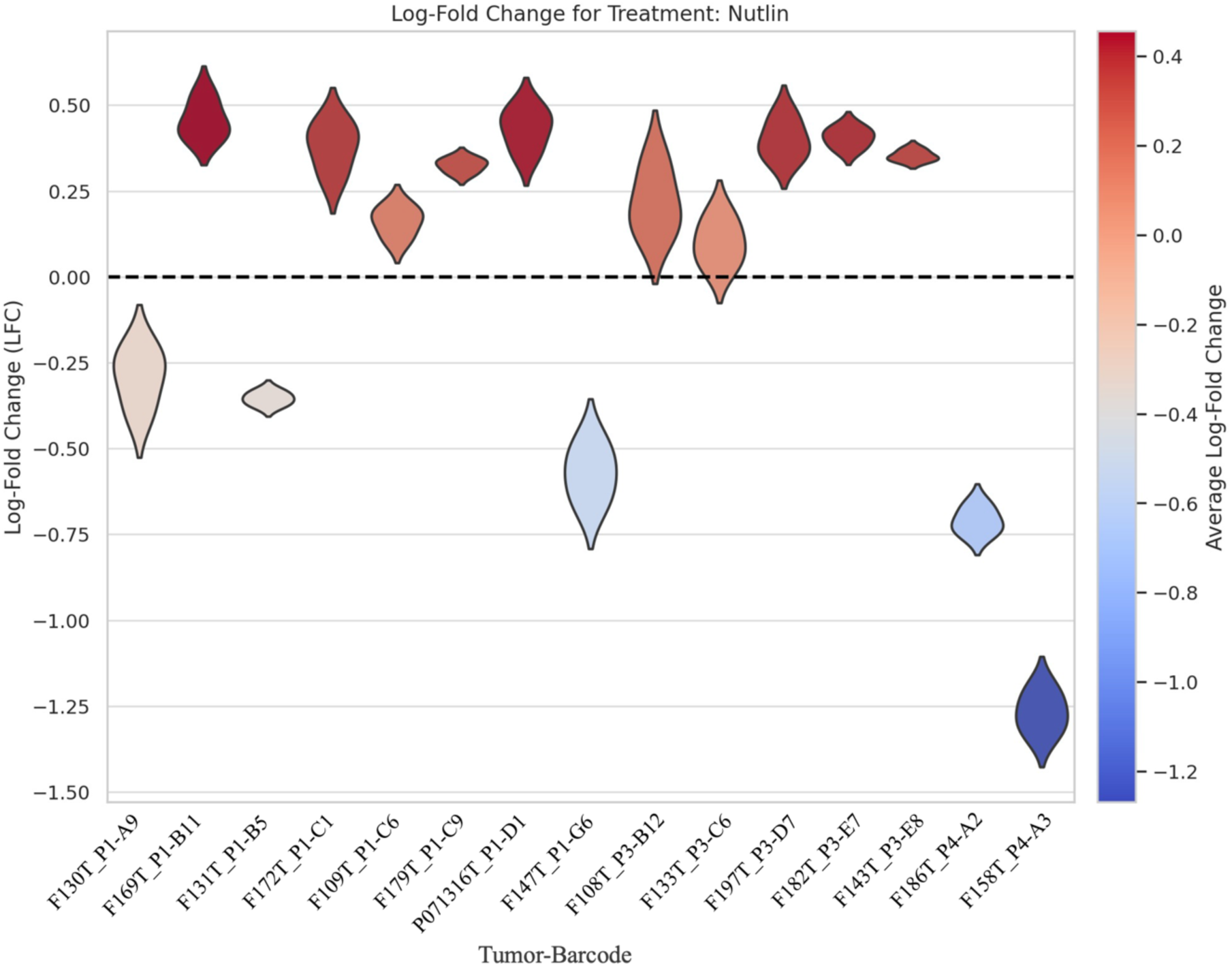
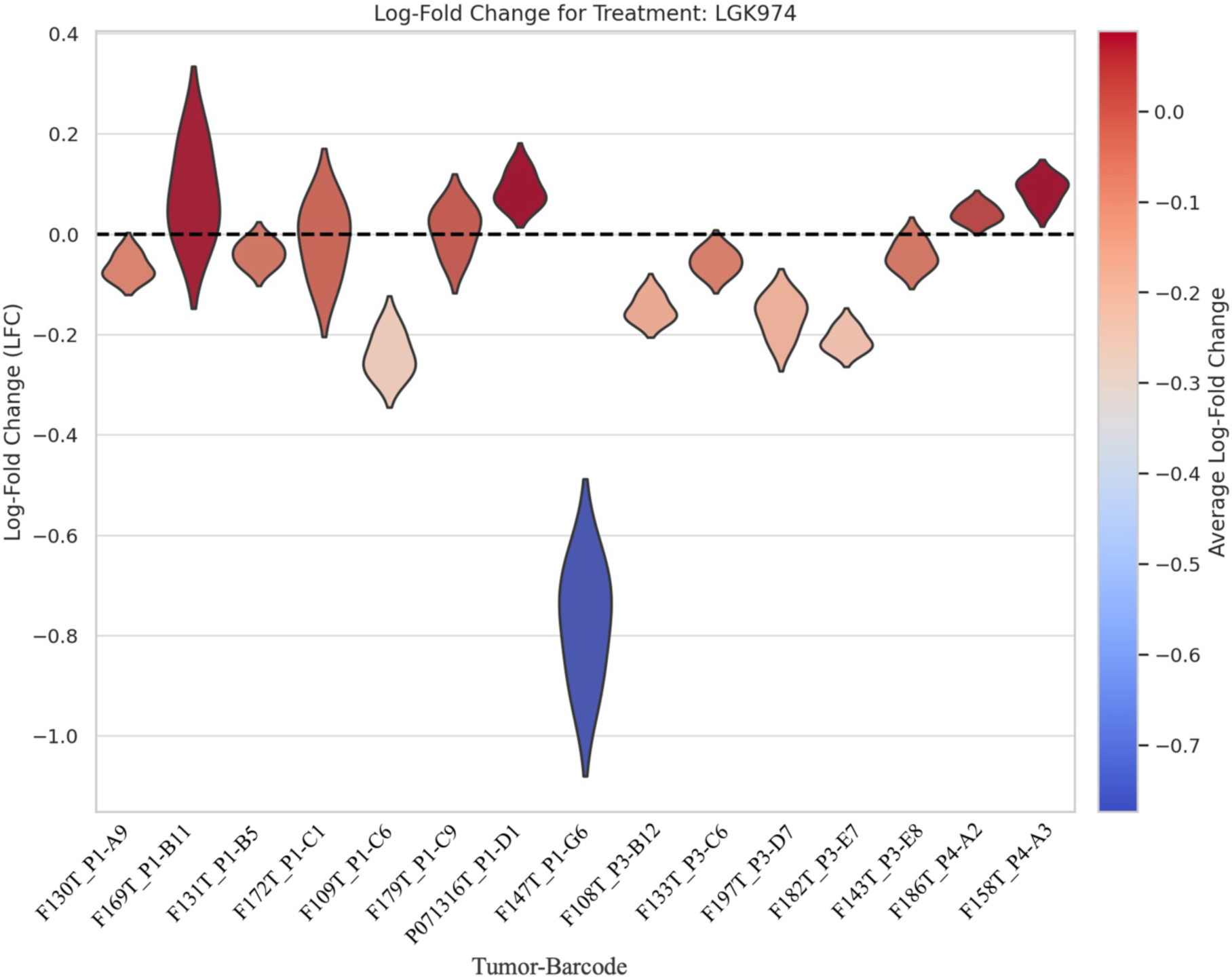
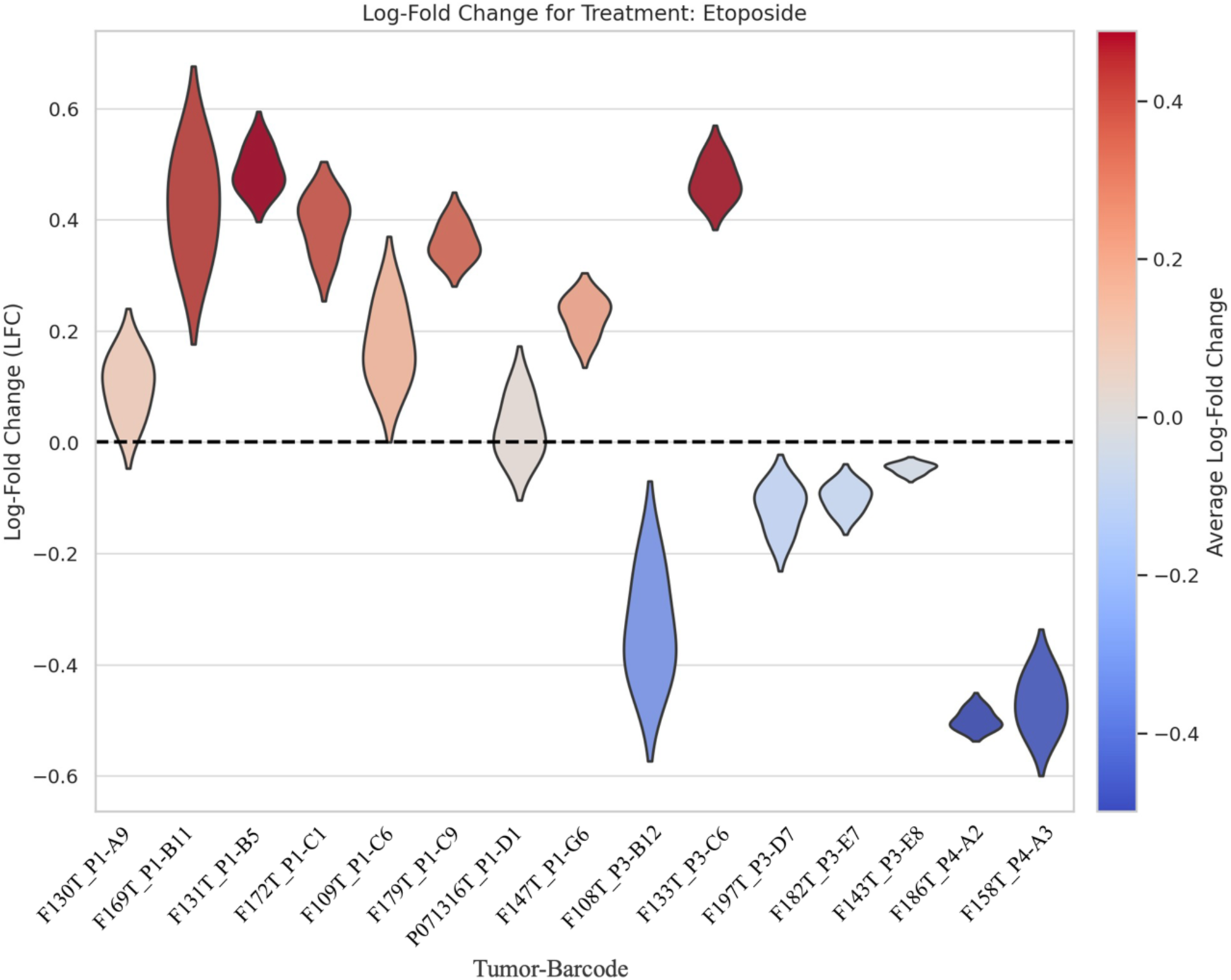
Drug responses of BC pool-2 PDOs over time, deconvoluted by NGS. Log-fold change in frequency of each barcoded PDOs within the pool-2, treated with different drugs for 27 days. Each violin represents the log-fold change in in a specific barcode relative to vehicle control.

In our previous study we discovered through a genome-wide CRISPR KO library screen that the F147T PDO had a PORCN gene knockout dependency, consistent with a role for self-stimulating (autocrine) WNT pathway activation in this RNF43-mutant tumor ^27^. In this study, we utilized LGK-974, a drug that targets the Wnt-specific acyltransferase porcupine (PORCN) and blocks maturation of endogenous WNT ligand ^28^. F147T_P1-G6 grown without external Wnt supplementation, demonstrated significant sensitivity to the PORCN inhibitor. This indicates that this tumor has the capability to activate the Wnt pathway using its own WNT ligand, thereby operating through an autocrine rather than paracrine mechanism ^29^. The data here shows the ability of this platform to detect the sensitivity of one RNF43 mutant colon tumor to LGK-974 within the pool using NGS.

### StarTrace, A Barcode qPCR Quantification Method

To augment NGS quantification of individual barcodes, we developed a rapid and cost-effective qPCR approach to quantifying barcodes within a mixture. We used a multi-color qPCR with common forward and reverse amplification primers and unique TaqMan probes targeting individual DNA barcodes. This allowed the rapid quantification of each barcoded PDO within the mixture and detect drug dependent effects on Darwinian fitness. To demonstrate the principal, we first created a 50/50 mixture of two BC PDOs, (F147T_P1-G6 and F130T_P1-A9) and treated the mixture with eight different concentrations of the PORCN inhibitors, LGK-974 and Wnt-C59. After the treatment was completed, we prepared genomic DNA and used StarTrace PCR to detect the two unique barcode sequences in the mixture and determine the quantity of each organoid in each condition. Figure 7 shows the relative sensitivities of F147T (harboring a somatic mutation to RNF43) and F130T_P1-A9. We next applied the StarTrace PCR assay to a pool of samples (Pool-2) treated with four different drugs. The results shown in figure 8, support the drug response pattern observed by NGS (Fig. 6). These findings demonstrate a robust correlation between NGS and StarTrace PCR method (Fig. 9), demonstrating the reliability of StarTrace primer/probe sets in accurately quantifying barcode signals in complex mixtures.

**Fig. 7:**
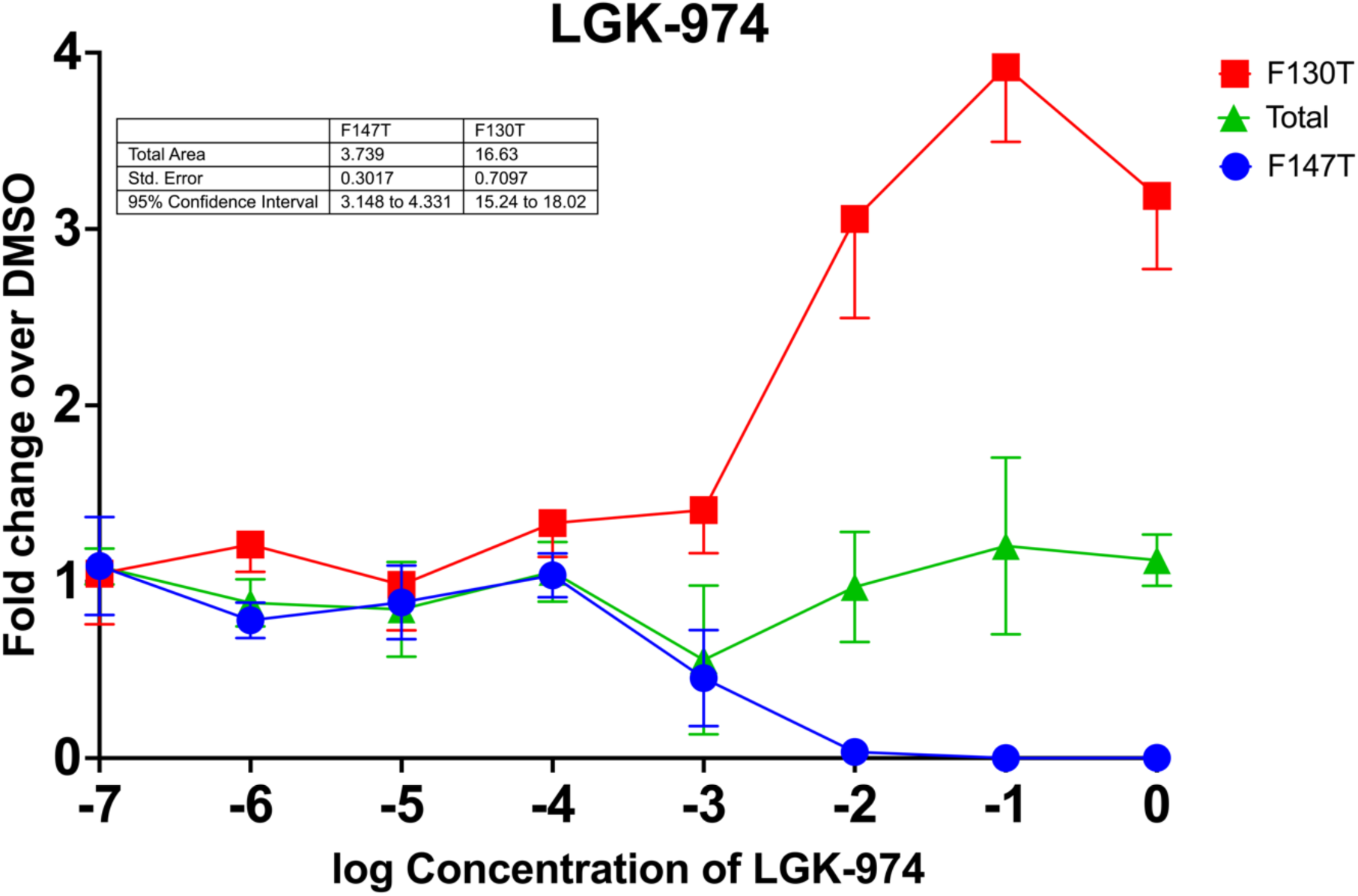
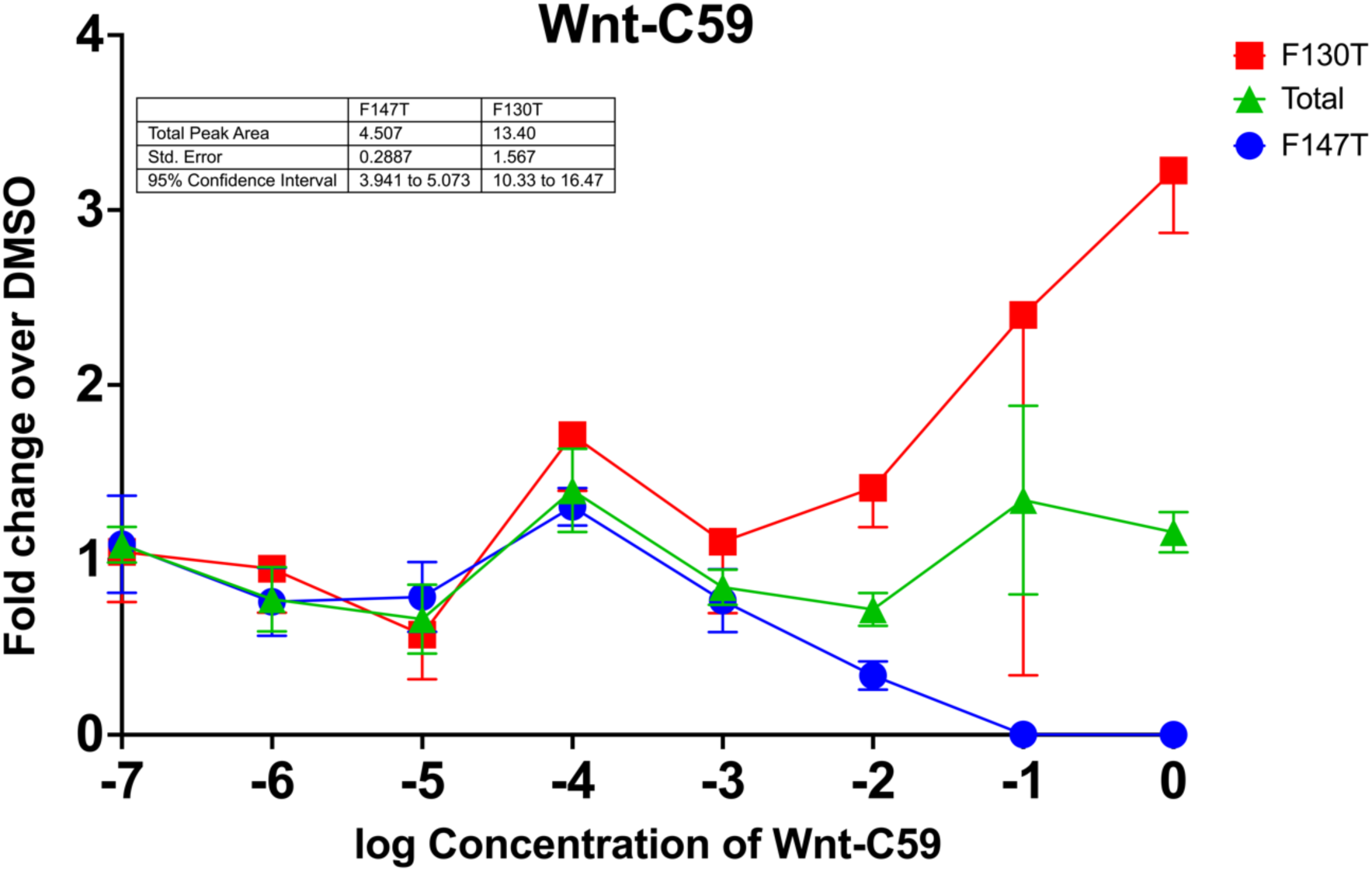
Dose response curve of PORCN inhibitors on a mixture of two BC PDOs, deconvoluted by StarTrace PCR. F147T_P1-G6 carried an RNF43 mutation and is sensitive to PORCN inhibitors (LGK-974 and wnt-C59). The ratios of the two tumors were measured by StarTrace PCR and plotted against the log concentration of LGK-974 and Wnt-C59.

**Fig. 8:**
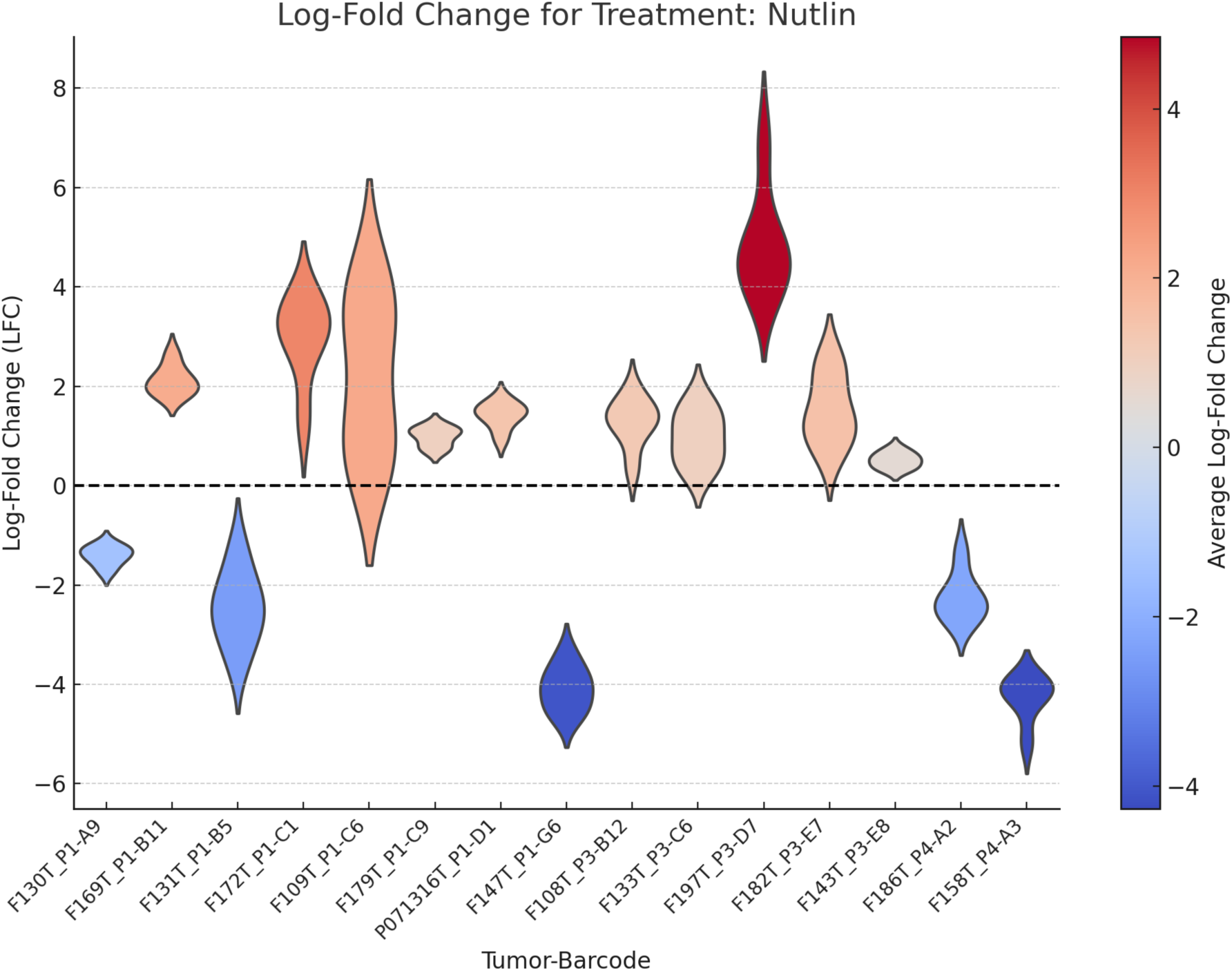
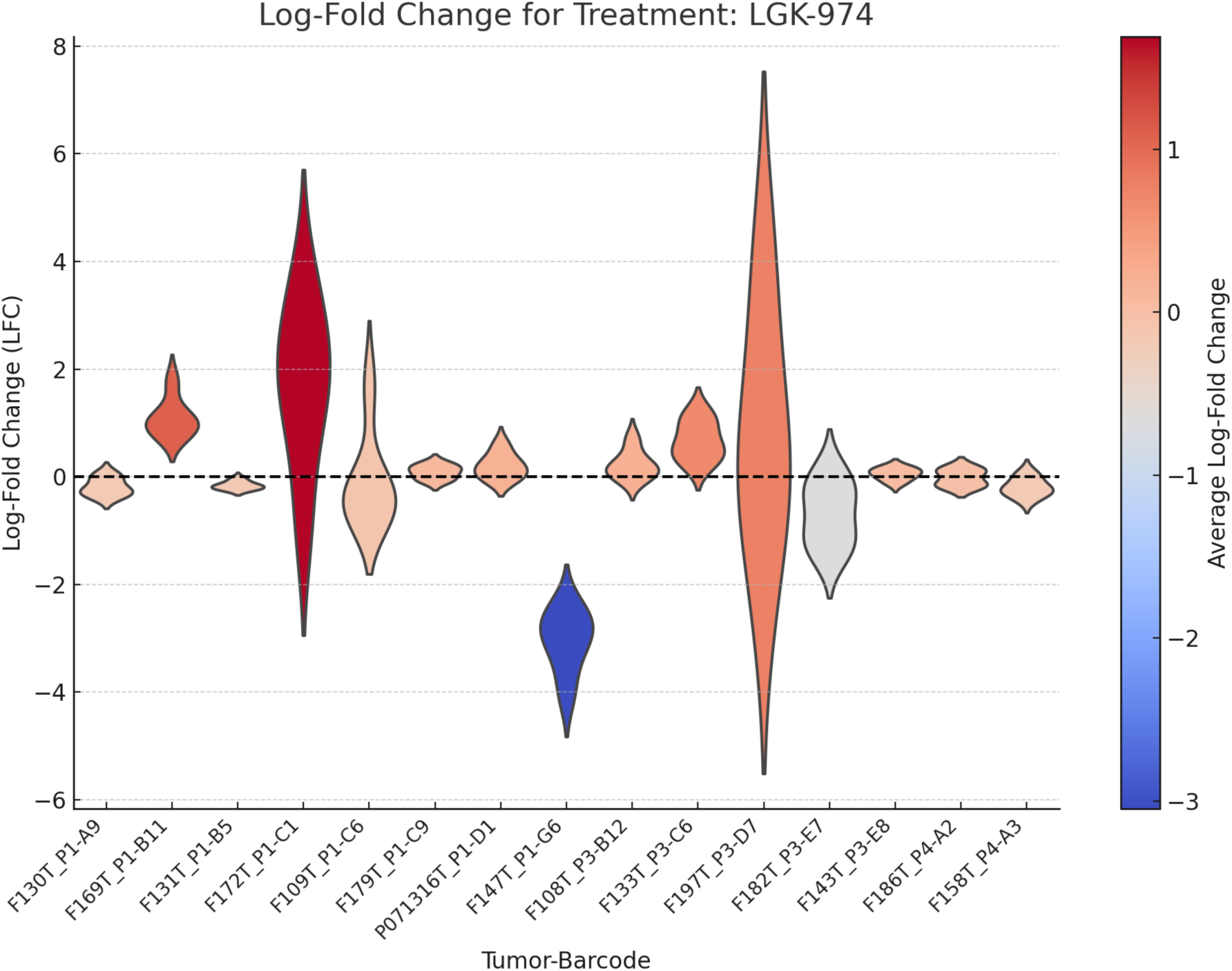
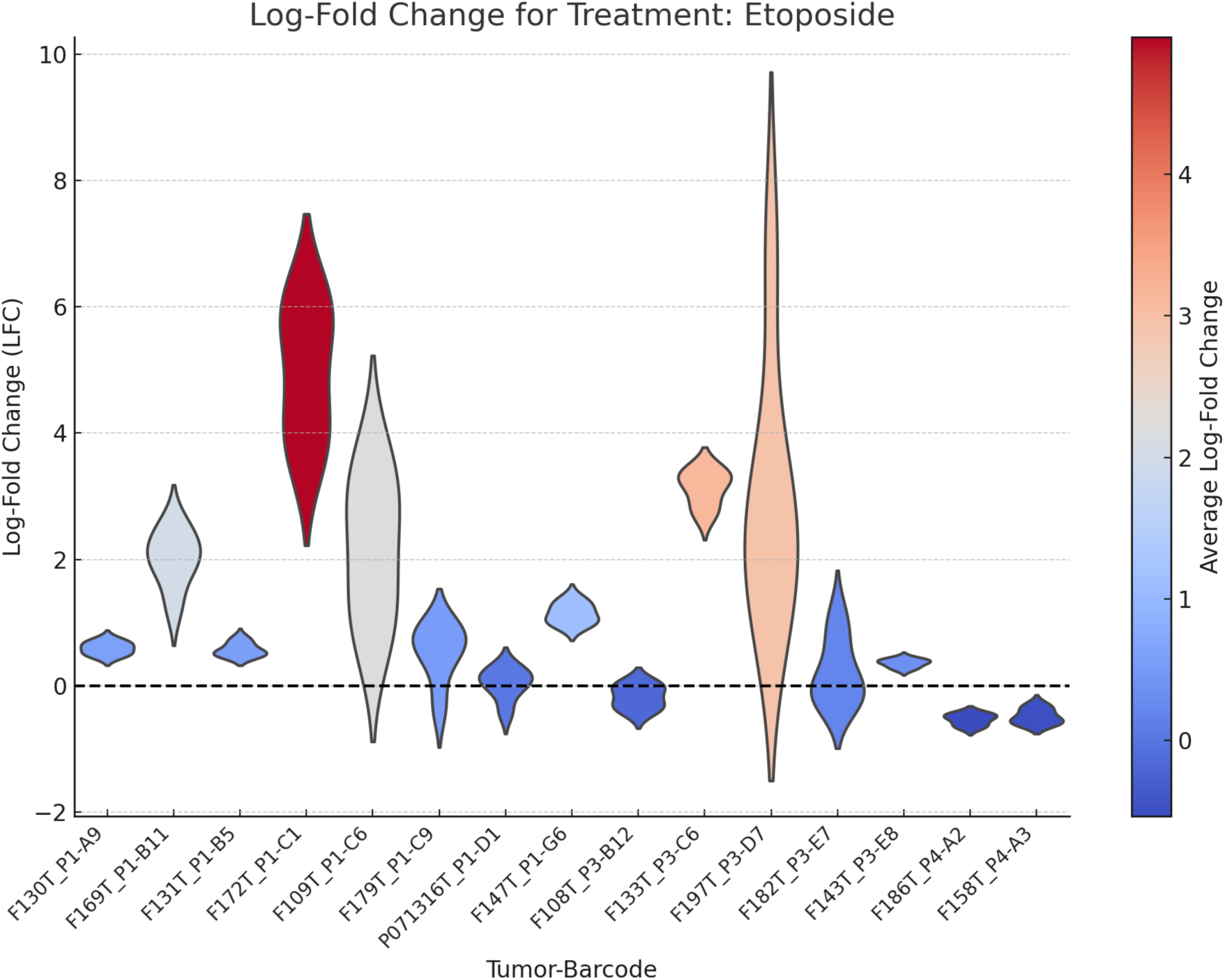
Drug responses of BC pool-2 PDOs over time, deconvoluted by StarTrace PCR. Log-fold change in frequency of each barcoded PDOs within the pool-2, treated with different drugs for 10 days. Each violin represents the log-fold change in in a specific barcode relative to vehicle control.

**Fig. 9:**
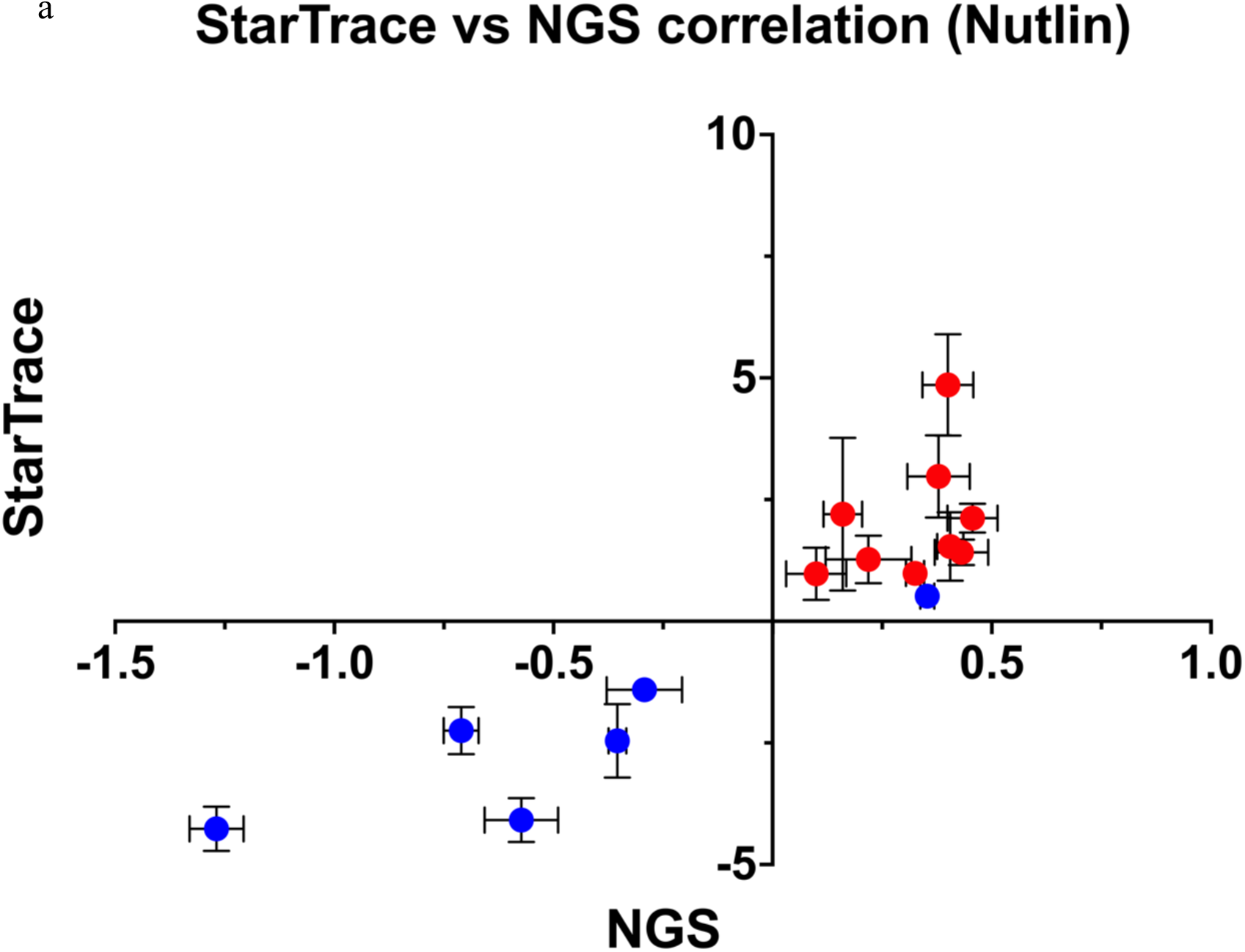
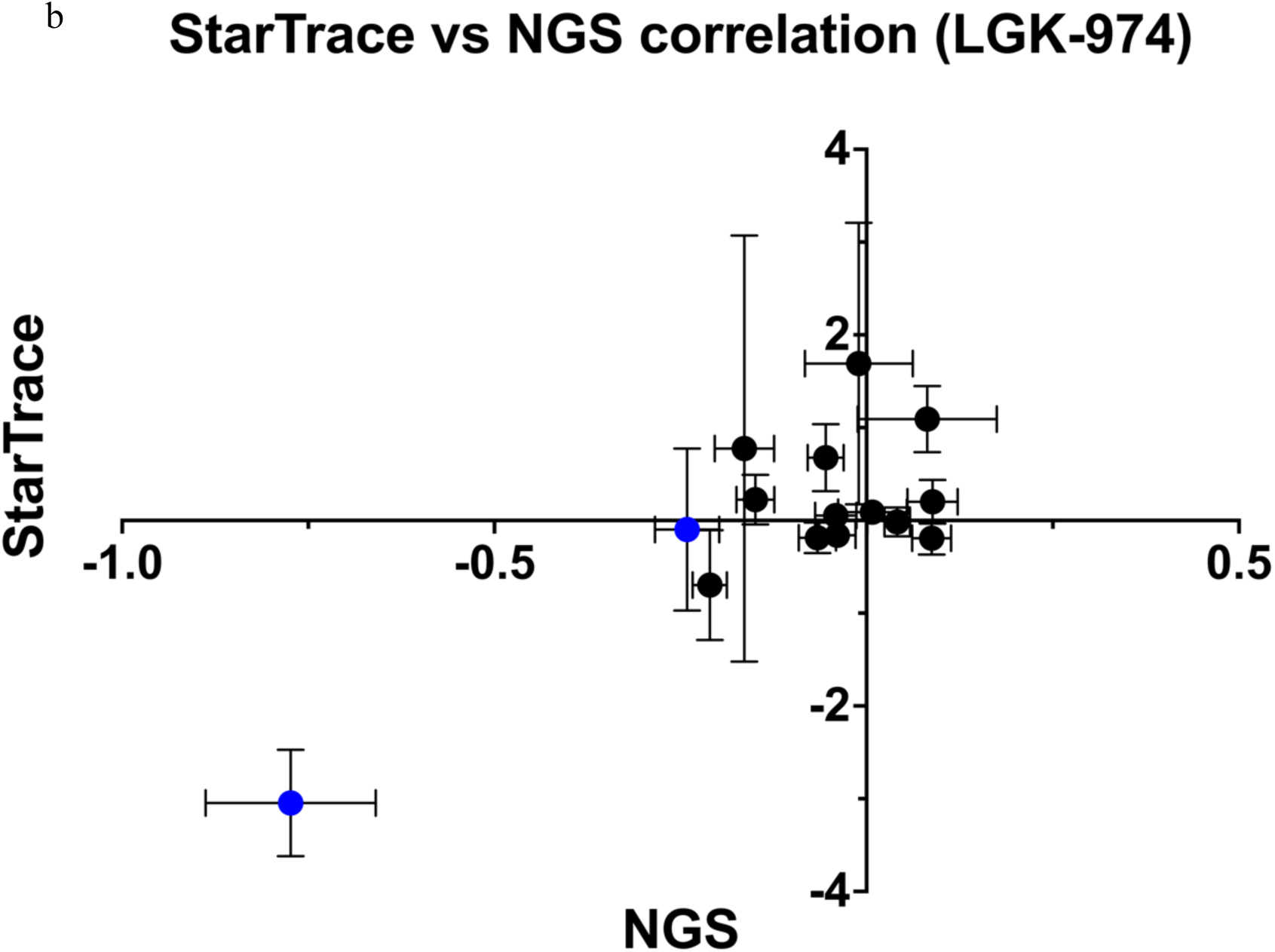
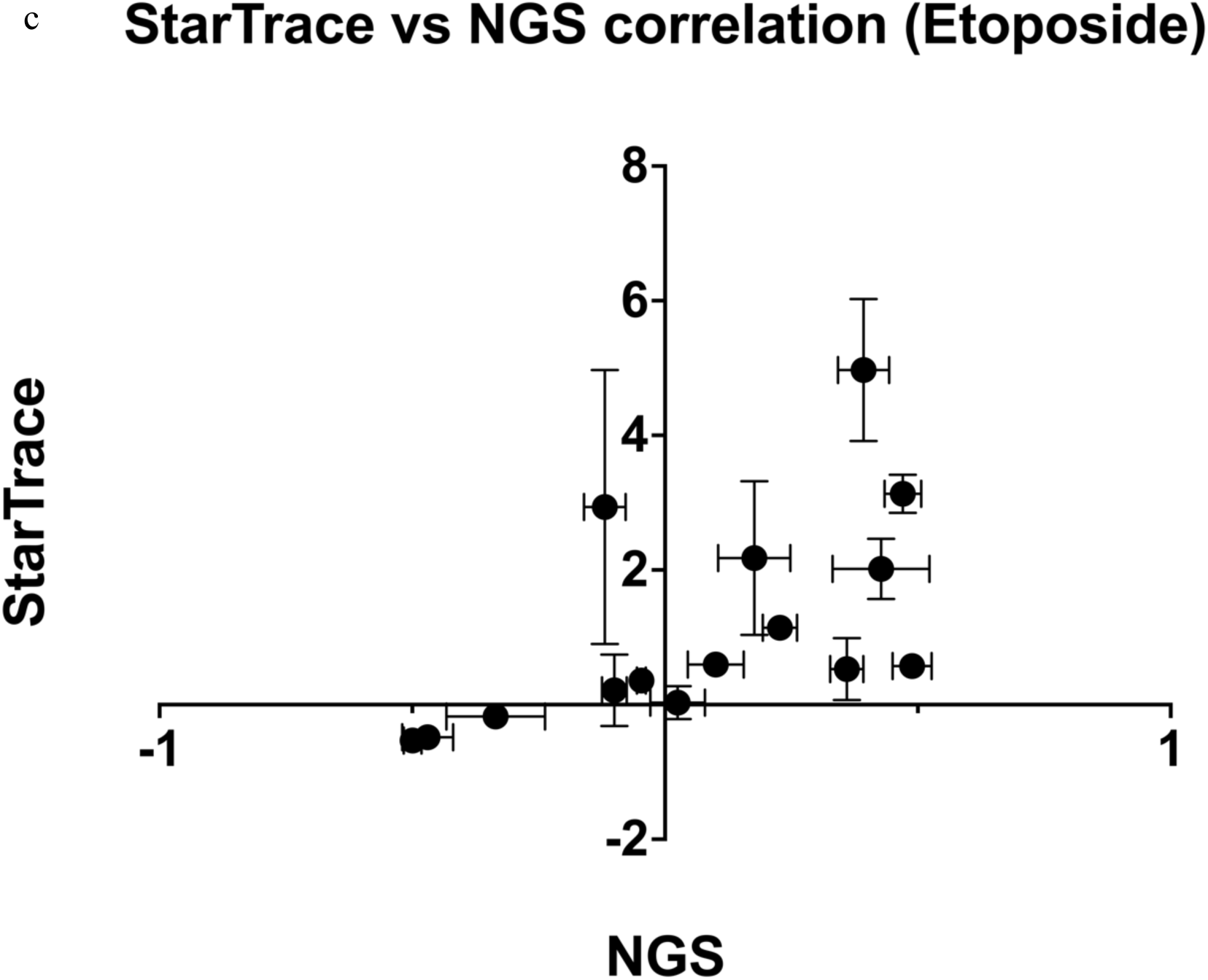
Correlation between NGS and StarTrace PCR. Comparison of the log-fold change in barcoded PDOs frequency obtained by NGS and StarTrace PCR for Nutlin (a) LGK-974 (b) and Etoposide (c). **a,** blue represents TP53-WT and red represents TP53-mutant. **b,** blue represents RNF43 mutant tumor. **c,** sensitivity and resistance to Etoposide.

We then miniaturized the StarTrace assay using Pool-2 barcoded PDOs. We applied a simple high-throughput organoid plating method in a 96-well plate format ^30^ and treated them with four different concentrations of 8 different drugs. In this miniaturized platform, we could successfully detect 12 of the 15 organoids in all time points of the untreated samples. Two organoids fell below the limit of qPCR detection. The heatmap in figure 10 shows the relative sensitivities of 12 out of 15 BC PDOs. The expected sensitivity to PORCN inhibitor was observed in the RNF43 mutated tumor (F147T_P1-G6). Interestingly, when this tumor was grown in the presence of external Wnt supplement, the efficacy of the PORCN inhibitor was notably diminished, aligning with the mechanism of PORCIN inhibitor action. StarTrace PCR correctly detected the expected nutlin sensitivity in the TP53-WT tumors F130T_P1-A9, F131_P1-B5, F147T_P1-G6, F186T_P4-A2, and F158T_P1-A3. These organoids’ nutlin sensitivities were independently confirmed through nutlin sensitivity assays on individual tumors grown separately and the results aligned with TP53 mutations obtained from nanopore-based TP53 amplicon sequencing and by Illumina-based whole exome sequencing. Sensitivity to Gefitinib, Etoposide, Daunorubicin, and Cisplatin were detected in select individual organoids, reflecting the diverse landscapes of therapeutic vulnerabilities within the sampled tumor populations. These findings underscore the potential for tailored therapeutics approaches based on specific drug sensitivity profiles observed in the organoids, highlighting the advantages of using this rapid, high-throughput, and precision assay.

**Fig. 10:**
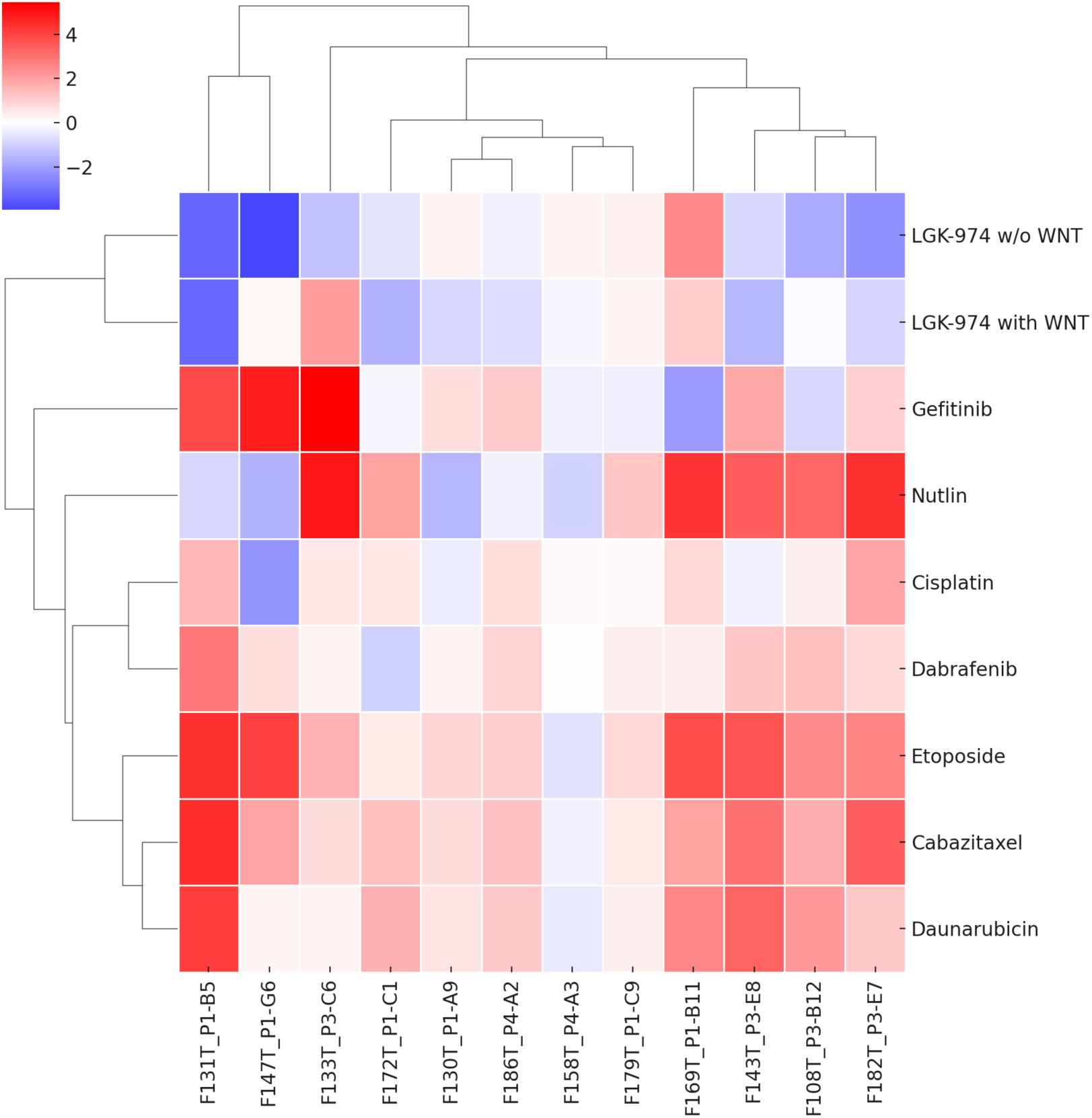
StarTrace PCR on the Pool-2 miniaturize assay. Clustered heatmap of the log fold changes in PDO frequencies over 10 days in presence of different drugs. The x-axis represents the various PDO samples. The y-axis represents the different drugs tested at the 10uM concentration. Negative log fold changed (cooler colors) indicate a reduction in cell numbers relative to untreated vehicle control (DMSO), suggesting dug sensitivity. Positive fold changes (warmer color) indicate an increase in cell numbers relative to untreated vehicle control (DMSO). 3 of the PDOs were not detectable so are not shown.

## Discussion

This study demonstrates the utility of PDO barcoding combined with molecular detection to assess drug sensitivity in mixed PDO populations. In some cases, drug sensitivity is mechanistically linked to known somatic mutations. Examples of this are TP53 mutations (13 PDOs) and Nutlin resistance (12 resistant PDOs), RNF43 mutations (2 PDOs) and sensitivity to PORCN inhibitors (1 sensitive PDO), and KRAS/BRAF mutations (6 PDOs) and resistance to lapatinib (6 resistant PDOs). The platform therefore accurately identified PDO sensitivity and resistance in 19 out of 21 cases (90%) where somatic mutations predicted drug response. This shows that the platform is useful for preclinical studies of novel drugs targeting known mutations.

In other cases, drug sensitivity or resistance are not understood in terms of somatic mutations. The platform still accurately identifies and quantifies drug response. The platform could be useful for making clinical choices for cancer patients by avoiding harsh chemotherapy regimens for which their tumors are resistant, and choosing treatments more likely to work based on positive results obtained from the organoid avatars. Future studies will be needed to determine the clinical predictive value of organoid avatar testing *ex vivo*.

The strong correlation between next-generation sequencing (NGS) and StarTrace PCR demonstrates that qPCR can often be used to detect subtle variations in drug response among different PDOs. The StarTrace PCR is relatively cost-effective, and accurate; however, it has limitations in sensitivity and scalability compared to sequencing-based detection. StarTrace encountered a tradeoff between achieving high sensitivity at detecting low levels of individual organoids and scaling the platform for multiple drug testing in miniaturized assay formats optimized for high throughput drug screening. Complementing StarTrace PCR with NGS can improve the detection of low-frequency cell populations and allow scaling up assays to enable high throughput drug screening. By leveraging both StarTrace PCR and NGS, we ensure precise tracking of treatment responses, facilitating tailored therapeutic strategies that accommodate the complex nature of individual tumors. This dual approach not only deepens our understanding of the genetic rules governing drug response, but also supports the strategic selection of personalized treatments, advancing the idea of personalized medicine. In conclusion, these findings show the value of PDOs in drug response assays and underscore the potential of combining advanced molecular techniques with traditional drug screening methods to enhance the precision of cancer treatment strategies. By integrating detailed genetic profiling with responsive organoid models, we can better predict therapeutic outcomes and tailor treatments to exploit the unique vulnerabilities of cancer, thereby improving patient outcomes in a clinical setting.

---

**Supplementary Fig. 1:**
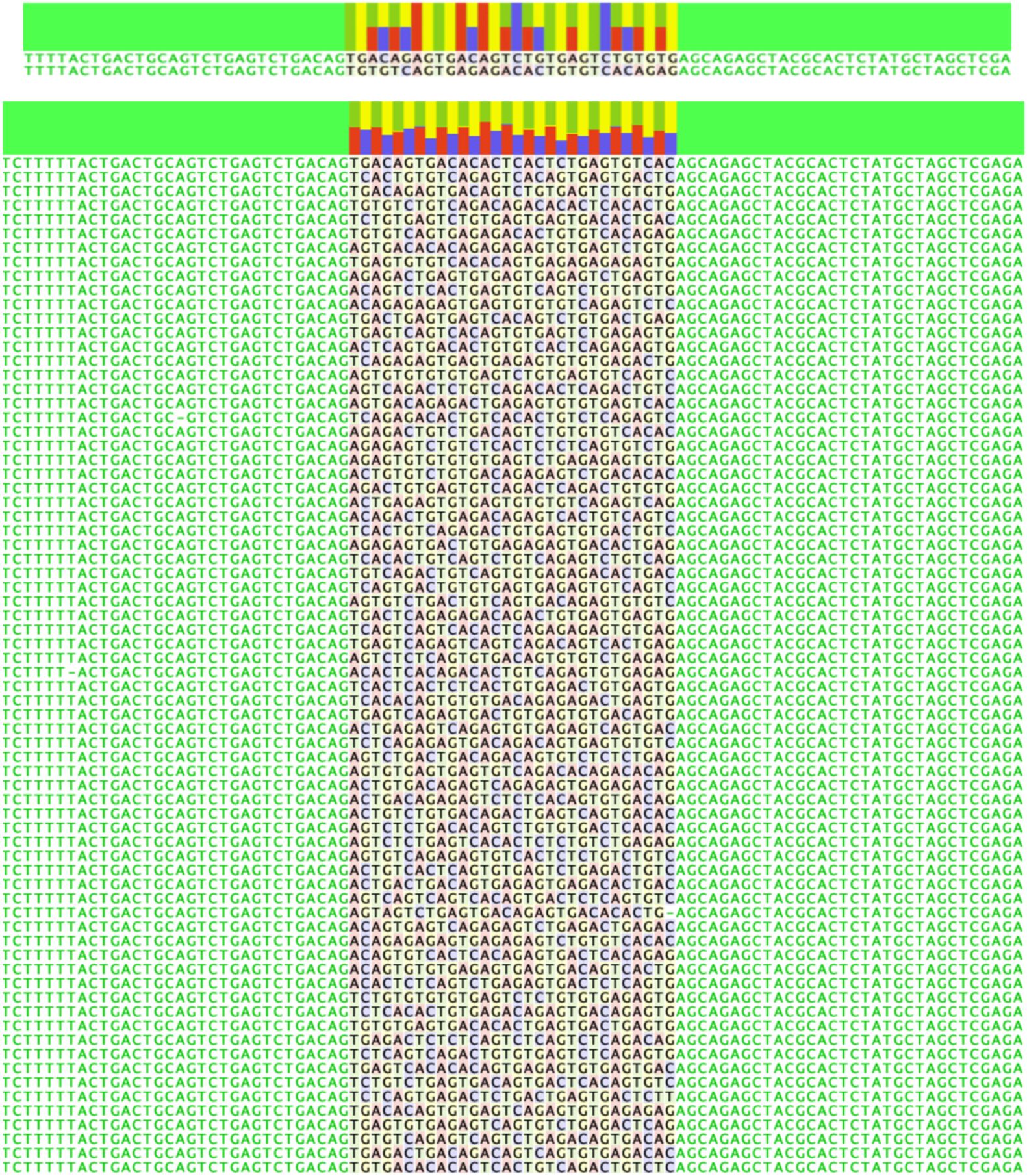
Sanger sequencing results and assembly of each 72 unique DNA barcode in relation to each other.

**Supplementary Fig. 2:**
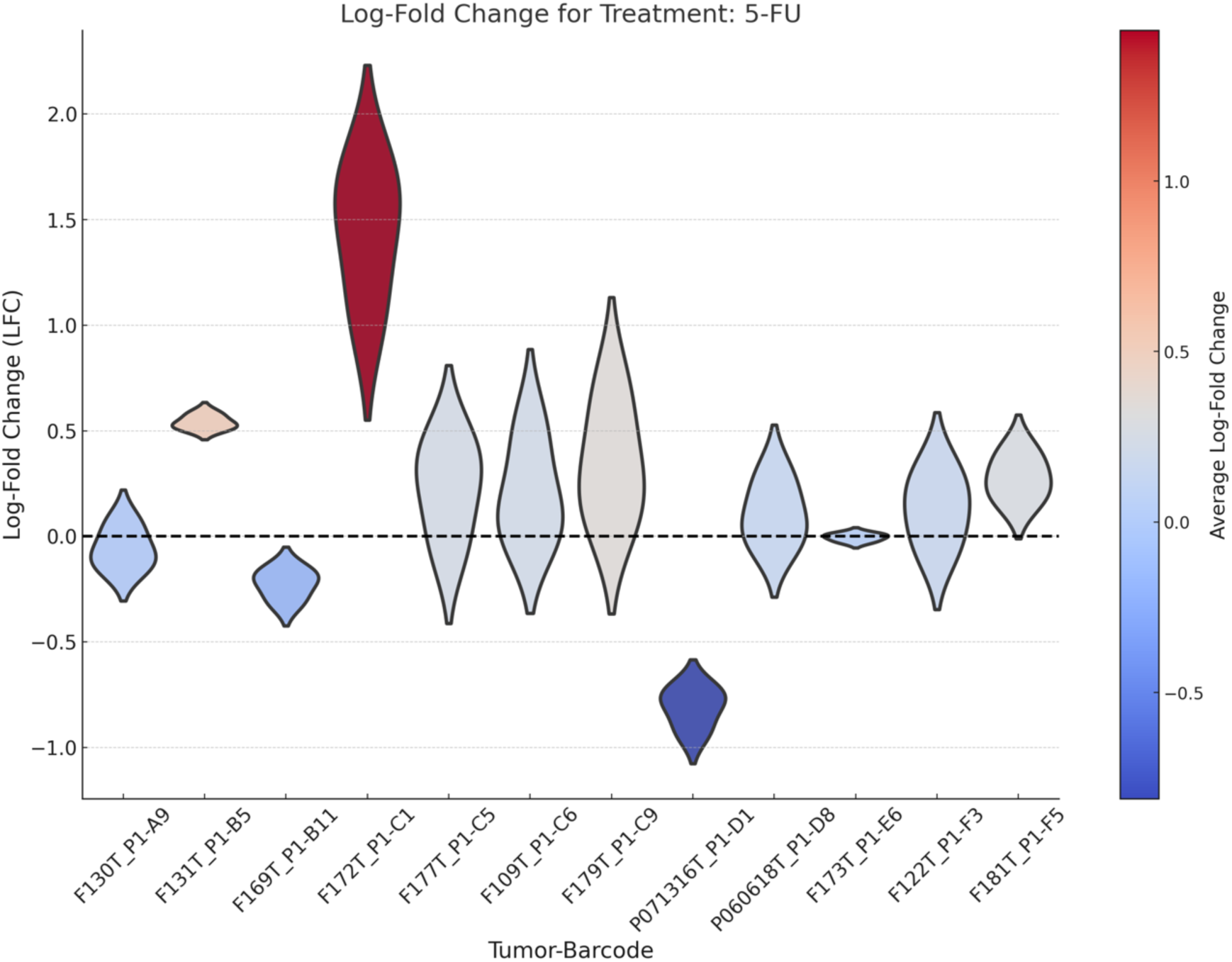
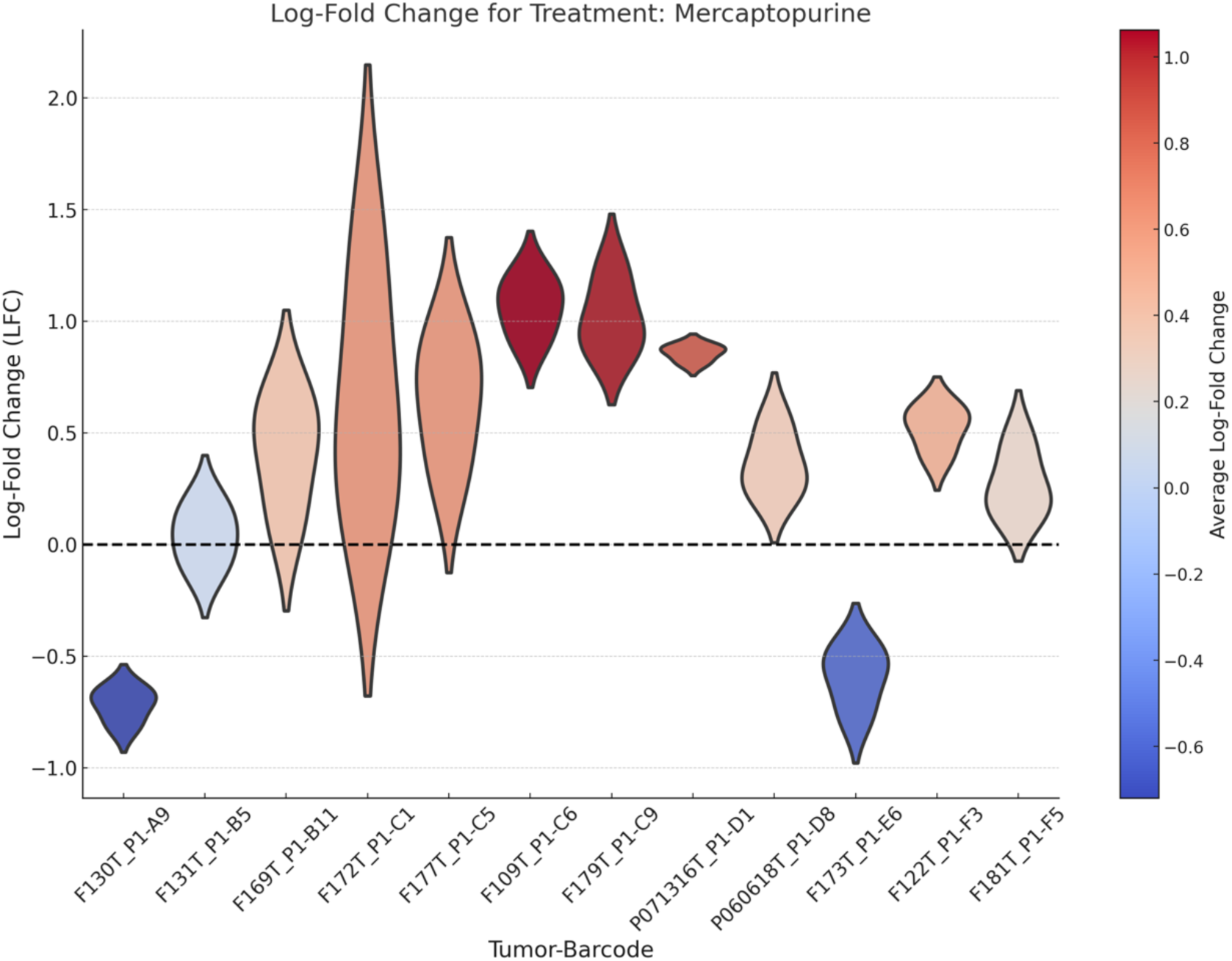
Drug responses of BC pool-1 PDOs over time, deconvoluted by NGS. Log-fold change in frequency of each barcoded PDOs within the pool-1, treated with different drugs for 28 days. Each violin represents the log-fold change in in a specific barcode relative to vehicle control.

## Methods

### Organoid Isolation and Culture

Organoids were created from surgically resected colorectal cancers ^31^. Small pieces (1cm square) of mucosal tissue were separated and washed in 1X PBS (Corning) supplemented with 0.1% Amphotericin B (Gibco) and 0.2% Primocin (Invivogen) to remove bacteria. Crypts were isolated with digestion and incubation in 1X PBS supplemented with 10% Collagenase A (Sigma) and 10 μM Y-27632 Rho Kinase inhibitor (Selleckchem) on a shaker at 37°C for 40 min, followed by addition of 5% FBS (Gibco) and incubation for 5 minutes at room temp. Crypts were dislodged mechanically from tissue by pipetting up and down, separated into a new tube for additional wash and were centrifuged at 500 x g for 5 min. The isolated stem cells were resuspended in 75ul of ice-cold RGF BME Matrigel (R&D systems) per well and plated in 12-well plates. The Matrigel was polymerized for 10-15 minutes at 37°C humidity-controlled incubator with 5% CO2 and subsequently covered with 2ml per well of human colon organoids media (ENRA) containing 10 μM Y-27632 RhoKinase inhibitor (Selleckchem. The ENRA media contained Advanced DMEM/F-12 media (Gibco) supplemented with 10% R-Spondin, 10% Noggin (all CM produced in-house), 1x N2, 1x B27, 10 mM HEPES, 1x Glutamax, 1% Penicillin/Streptomycin, 0.25 ugmL−1 Amphotericin B, 50 ngmL−1 human Epidermal Growth Factor EGF (all from Gibco), 10 mM Nicotinamide, 1.25 mM N-acetyl-cysteine, 10 nM Gastrin (all from Sigma), 0.5 μM TGF-β type I receptor inhibitor A83-01 (Tocris Bioscience), 100 ugmL−1 Primocin (Invivogen). Media was changed every other day. For the first two changes of media after culturing, media contained 10 μM Y-27632 RhoKinase inhibitor (Selleckchem). Organoids were passaged 1:4 every 10-14 days. For passaging, organoids and Matrigel were covered with 2ml/well of TrypLE (Gibco) plus 10 μM ROCK inhibitor (Selleckchem), dissociated by incubation at 37°C for 5-7 min, then mechanically disrupted using a P1000 pipette and transferred into a 15-ml conical tube. Centrifuged at 500 x g for 5 min. The pellet was resuspended and plated in ice-cold fresh Matrigel (R&D systems) and covered with ENRA media. Organoid lines were constantly tested for mycoplasma contamination and resulted negative.

### Cell Culture

All colon tumor organoids were grown as described before ^31^. Organoids were maintained in ENRA media containing 1% Penicillin/Streptomycin, 100 μg/ml Primocin, 0.25 ug/ml Amphotericin B. Organoids were kept in a 37°C humidity-controlled incubator with 5% CO2 and were maintained in exponential phase growth by passaging every 10-14 days and media was changed every other day. For viral transduction, 6μg/ml polybrene was added. After viral transduction, 2μg/ml puromycin were used for selection.

Organoid mixtures were treated with sub-lethal concentrations of each drug over a span of 28 days or 27 days with media changes and drug replenishment every other day, and the organoid pool replated in fresh Matrigel every week.

### Isolation Individual Clone Barcode from the Barcoding Pool

ClonTracer Barcoding pool Library DNA was purchased from addgene (#67267). DNA was transformed into the bacteria and individual colonies were picked. Plasmids were purified from each individual clone and confirmed by sanger sequencing. Lentivirus of 72 individual unique barcodes were produced by the Duke University Viral Vector Core facility.

### Barcoding Patient’s Derived Organoids - Viral Transduction

Prior to viral transduction, organoids were digested by incubation in TrypLE contained 10 μM ROCK inhibitor at 37°C for 10-20 min. Then, cells were resuspended in 1X human colon tumor organoids media (ENRA) contained 10μM ROCKi and Polybrene (6μg/ml). Next, viral sufficient to achieve a multiplicity of infection (MOI) of approximately 3, were added to the cell mixture. The mixture was then transferred to a single well of a 24-well tissue culture plate, which was spun at 1000 x g for 2 h at 30°C. After centrifugation, the plate was incubated at 37°C incubator with 5% CO2 for additional 3 h (total of 5 h). Following the incubation period, virus was washed out from the organoids with ice-cold PBS contained 10μM ROCKi. Transduced organoids then were embedded in ice-cold Matrigel and covered with ENRA media containing 10μM ROCK inhibitor for the first two media changes (4 days). For all the other feedings, the organoids were maintained in ENRA media without ROCKi. The Puromycin selection was started 2 days after the viral transduction and continued for 7 days.

### Next-generation Sequencing (NGS) Assay

First, equal cell numbers of each barcoded organoids were mixed. Organoid mixtures were either plated in domes of Matrigel in tissue culture plates (48-wells, 12-wells or 6-well plates). A total of ∼250,000 cells per replicate were plated for drug treatment experiments. Organoid mixtures were treated with sub-lethal concentrations of each drug over a span of 28 or 27 days with media changes and drug replenishment every other day, and the organoid pool replated in fresh Matrigel every week. At weekly time points, 50% of the organoid pool was removed for sequencing and the remaining 50% was replated in fresh Matrigel. Throughout sampling for DNA purification, barcode PCR and amplicon sequencing, the size of each barcoded PDO population never dropped below a 1,000-fold bottleneck of 15,000 cells.

The genomic DNA of pooled barcoded PDOs were PCR amplified with primers that flank the DNA barcode sequences (Barcode_backbone_F and Barcode_backbone_R) and the products sequenced by Oxford Nanopore platform (LSK109 and NBD112.24 (Q20+)).

Barcode primers used for DNA barcode sequence amplification:

Barcode_backbone_forward primer, 5′–3′; CGATTAGTGAACGGATCTCGAC

Barcode_backbone_reverse primer, 5′–3′; CCATTTGTCTCGAGCTAGCATA

Primers used for ERBB3 amplification:

ERBB3_exon3_Forward primer, 5′–3′; TTGCCCTGTTGTCTCTCTCA

ERBB3_exon3_Reverse primer, 5′–3′; GTGGCTGGAGTTGGTGTTAT

### StarTrace PCR

Equal cell numbers of each barcoded organoids were mixed for pool-2. They were resuspended in a slurry of ENRA:Matrigel (3:4 ratio) and plated in rings around the rim of each well of a 96-well tissue culture plate (∼10,000 total cells per well) as described ^30^. Select drugs from the approved Oncology Drug set VI were screened at maximum concentration of 10uM and four serial dilutions over span of 10 days with refeeding every other day. After the treatment was completed, a multi-color fluorescent probe-based real time PCR assay (Taqman) was performed to quantify each barcoded organoids within the mixture. The primers targeted the barcode sequence location, and each probe are designed in a way to detect one unique barcode sequence which allowed us to quantify each unique barcoded organoids within a mixture and detect the effect of any drug concentration on each one separately. A probe targeting common backbone sequence was used to internally normalize. The assay used 4μL of 5x PerfeCTa Multiplex qPCR ToughMix (Quantabio), 6μL of triplex Primers and Probes mix, gDNA as template and water to adjust the reaction volume to 20μL. PCR cycling conditions: 1min at 95°C for 1 cycle; 10s at 95°C, 30s at 62°C for 4 cycles; 10s at 95°C, 30s at 59°C for 4 cycles; 10s at 95°C, 30s at 56°C for 4 cycles; association curve from 95°C to 55°C.

primers and probes used for qPCR assay:

Barcode_forward primer, 5′–3′; CGATTAGTGAACGGATCTCGAC

Barcode_reverse primer, 5′–3′; CCATTTGTCTCGAGCTAGCATA

Barcode_backbone_probe; /5TexRd-XN/ACTGACTGCAGTCTGAGTCTGACA/3IAbRQSp/

P1-A9_probe; /5HEX/AGTGACTGT/ZEN/GTGAGTGAGAGTGTCAGT/3IABkFQ/

P1-B5_ probe; /5Cy5/TGACAGAGA/TAO/GTCTCTCACAGTGTGACA/3IAbRQSp/

P1-B11_ probe; /56-FAM/ACTGTCTGT/ZEN/GACAGACTGAGTCAGTGA/3IABkFQ/

P1-C6_ probe; /5Cy5/ACAGTGAGT/TAO/CAGAGAGTCTGAGACTGAG/3IAbRQSp/

P1-C9_ probe; /5HEX/AGACTGTGA/ZEN/GTGTCAGACTCAGACTGT/3IABkFQ/

P1-D1_ probe; /5Cy5/TGAGACTCT/TAO/CTCAGTCTCAGTCTCAGACA/3IAbRQSp/

P1-G6_probe; /56-FAM/TGTCACTCA/ZEN/GTGTGAGTCTGAGACTGT/3IABkFQ/

P3-B12_ probe; /56-FAM/ACAGTCTGA/ZEN/CAGTGAGTGTGTGTCAC/3IABkFQ/

P3-C6_ probe; /56-FAM/TCTGTGAGT/ZEN/CTGTGAGTGAGTGACACT/3IABkFQ/

P3-D7_ probe; /5Cy5/AGTGTCTGA/TAO/CTGTCAGTGACAGAGTGT/3IAbRQSp/

P3-E7_ probe; /5HEX/TGTCAGAGT/ZEN/CAGTCTGAGACAGTGACA/3IABkFQ/

P3-E8_ probe; /56-FAM/ACAGTGTGT/ZEN/GACAGAGAGACTGAGT/3IABkFQ/

P4-A2_ probe; /5HEX/AGTGACTGT/ZEN/CTGACTGACTCTGACTCAC/3IABkFQ/

P4-A3_ probe; /5Cy5/AGTCTCTCA/TAO/GTGTGACAGTGTGTCTGA/3IAbRQSp/

The following 10x primer and probe mix was made for each pair:

5μM forward primer; 5μM reverse primer; 2.5μM probe

A total of 5 different mixtures, each containing an equal amount of 4 different primers and probe, was prepared as follow:

| Mixtures | Barcode_FAM | Barcode_HEX | Barcode_Cy5 | Barcode_TEXAS RED |
| --- | --- | --- | --- | --- |
| 1 | F147T_P1-G6 | F130T_P1-A9 | F158T_P4-A3 | Barcode_Backbone |
| 2 | F143T_P3-E8 | F186T_P4-A2 | F131T_P1-B5 | Barcode_Backbone |
| 3 | F108T_P3-B12 | F182T_P3-E7 | F197T_P3-D7 | Barcode_Backbone |
| 4 | F133T_P3-C6 | F172T_P1-C1 | F109T_P1-C6 | Barcode_Backbone |
| 5 | F169T_P1-B11 | F179T_P1-C9 | P071316T_P1-D1 | Barcode_Backbone |

### Genomic DNA isolation

Genomic DNA (gDNA) was isolated using DNAdvanced kit (BeckmanCulter). The gDNA concentrations were quantitated by Qubit.

### NGS Analysis

Raw FASTQ files were analyzed by a fuzzy text matching program (UGREP). Example of the command:

ugrep -Z3 -c ’guide_forward_seq|guide_reverse-complement_seq’ sample_fastq_file.fastq

-Z3 is the fuzzy option to allow 3 mismatches.

Each FASTQ file from a single sample replicate was queried for all expected barcodes and the resulting raw counts normalized to the sum of all counts. Logfold changes relative to the no-drug control was used to assess drug-dependent effects on Darwinian fitness.

### StarTrace Analysis

For every well, we first calculated the difference between the Ct signal of each BC PDO and the TexasRed Ct (Backbone) signal. We then calculated the log fold change (LFC) of each BC PDO’s frequency in drug-treated samples, normalized to DMSO. A negative value means fewer total cells in the drug-treated samples compared to DMSO.

## Acknowledgment

Organoid biobanking was supported by an NCI SBIR to Dr. Carolyn Banister.

## Ethics Statement

All samples were collected under an IRB approval protocol (Pro00022064) “The Palmetto Health – University of South Carolina Biorepository”.

## Author Contributions

S.K and P.J.B conceptualized the StarTrace PCR method. S.P and P.J.B designed and optimized the NGS barcode detection assay. S.K, S.P, and P.J.B designed all the other experiments. S.K, S.P, N.P, V.M, S.L, E.G, R.B carried out all experimental work. S.K, S.P, N.P, and P.J.B designed and carried out all computational analysis. S.K, S.P, and C.E.B generated the barcoded patient’s organoids. C.E.B established colon human tumor’s organoid biobank. S.E.M provided colon human tumor tissues. P.J.B supervised the work. S.K and P.J.B wrote the manuscript, with input from all the authors.

## Competing Interests

Sana Khalili, Phillip Buckhaults, Shrey Patel, Carolyn Banister, and University of South Carolina have filed U.S. Patent Application No. 63/721,702 (StarTrace: A Multiplex Drug Testing Platform for Personalized Medicine), for aspects of this work on November 18, 2024. The other authors declare no competing interests.

## Supplementary Information

Supplementary information is available for this manuscript.

Correspondence and requests for materials should be addressed to

## Supplementary Information

## Supplementary Data.1

Somatic mutations in driver genes in all organoids. Patient-matched normal organoids were used as control.

## Supplementary Data.2

Barcode sequences and the sequencing raw counts used for analysis and tracking treatment responses.

